# Sequence context decodes multi-state conformational heterogeneity from crystallographic B-factors

**DOI:** 10.64898/2025.12.05.692009

**Authors:** Yiquan Wang, Minnuo Cai, Yahui Ma, Tieliu Shi, Kai Wei

## Abstract

Protein function is governed by conformational ensembles, yet the overwhelming majority of experimentally determined structures capture only static snapshots, leaving the multi-state heterogeneity that underlies biological activity largely inaccessible. X-ray crystallography, the dominant source of these structures, does carry one experimental proxy for motion, the atomic B-factor, but crystal packing and refinement artifacts frequently obscure its link to solution-state dynamics. Here we show that sequence context is sufficient to decode a protein’s multi-state conformational heterogeneity from this seemingly noisy signal. We introduce the B-Factor Corrector (BFC), a fine-tuned protein language model that reframes B-factor analysis from a purely physical calculation into a sequence-context-aware translation task. By leveraging deep contextual embeddings to decouple intrinsic conformational heterogeneity from lattice constraints, BFC recovers the residue-wise heterogeneity ranking derived from crystallographic structural ensembles, reproducing the normalized profile at a Pearson correlation of 0.83 while correcting the distortions that analytical formulas leave in the raw signal. We further demonstrate the utility of this sequence-inferred signal across structural-biology applications: it provides an integrable prior for molecular docking that is statistically equivalent to experimental B-factors, unmasks cryptic conformational heterogeneity essential for chaperone function, and guides the energetic refinement of antibody interfaces to resolve steric conflicts. These results suggest that the static structures of the Protein Data Bank can be effectively repurposed to decode the conformational heterogeneity latent within them.

## Introduction

Protein function is fundamentally dynamic, relying on conformational ensembles that extend beyond static coordinates (Frauenfelder et al., 1991; Csermely et al., 2010; Nussinov et al., 2023; Fenwick et al., 2014). While X-ray crystallography has contributed over 190,000 structures to the Protein Data Bank (PDB) (Berman et al., 2000), these models are traditionally viewed as static snapshots (DePristo et al., 2004). The atomic displacement parameter, or B-factor, provides the primary experimental metric for motion; however, its interpretation remains challenging. Influenced by non-biological factors such as crystal packing contacts, solvent conditions, and refinement heterogeneity, B-factors are frequently regarded as qualitative indicators of local disorder rather than quantitative measures of solution-state dynamics (Li and Brüschweiler, 2009; Carugo, 2022; Klyshko et al., 2024; Eyal et al., 2005).

To bridge the gap between static structures and dynamic function, researchers typically rely on molecular dynamics (MD) simulations (Karplus and McCammon, 2002) or Nuclear Magnetic Resonance (NMR) spectroscopy (Lasorsa and van Der Wel, 2025). While accurate, these methods are computationally expensive or experimentally demanding, leaving the vast majority of PDB entries without quantitative dynamic profiling. Elastic Network Models (ENMs) such as Gaussian Network Models (GNM) (Bahar et al., 1997) and Anisotropic Network Models (ANM) (Atilgan et al., 2001) offer rapid approximations but, as they rely on coarse-grained contact topology while treating amino acids as uniform mechanical nodes, often overlook the subtle chemical specificity governed by the local sequence environment.

Recent advances in deep learning (Xu et al., 2026; Wang et al., 2025a) have demonstrated that protein sequences inherently encode dynamic information accessible without explicit physical simulation. Early methods such as DynaMine (Cilia et al., 2014) predicted backbone dynamics directly from sequence, while tools like SETH (Ilzhöfer et al., 2022) and flDPnn (Hu et al., 2021) leveraged protein language models or composite features to predict intrinsic disorder propensities. More recently, sequence-based models have moved toward direct prediction of physical flexibility metrics. Pandey et al. (2023) showed that crystallographic B-factors can be largely predicted from primary sequence alone, OPUS-BFactor (Yang et al., 2025) integrated ESM-2 embeddings with structural features for B-factor prediction, and PEGASUS (Vander Meersche et al., 2025b) achieved prediction of MD-derived Root Mean Square Fluctuation (RMSF) from sequence using multiple protein language model representations. Structure-based approaches such as RMSF-net (Song et al., 2024) further demonstrated that integrating atomic coordinates with cryogenic electron microscopy (cryo-EM) density improves RMSF prediction accuracy. Pushing this further, SeqDance and ESMDance embed molecular-dynamics- and normal-mode-derived flexibility directly into a protein language model, predicting per-residue fluctuation and residue comovement from sequence alone and transferring this dynamic signal to downstream tasks such as mutation-effect prediction (Hou et al., 2026). While these tools collectively confirm that sequence and structure encode exploitable flexibility signals, they uniformly operate as *predictors*, mapping inputs directly to flexibility outputs. Where these tools predict flexibility *de novo*, BFC operates as a *corrector* that leverages sequence context to denoise existing experimental B-factor measurements, decoding the intrinsic conformational-heterogeneity signal from the crystallographic artifacts that obscure it.

We postulate that the discrepancy between crystallographic B-factors and solution-state dynamics represents a systematic artifact encoded in the local physicochemical context rather than random noise. This hypothesis is supported by preliminary validations in which experimental B-factors correlate significantly with MD-derived, single-trajectory RMSF across structurally diverse folds (Wang et al., 2025b), confirming that B-factors track genuine thermal dynamics; the systematic gap we target here is instead their weak correspondence with the cross-deposition ensemble heterogeneity that no single trajectory captures. Protein sequences likely store an evolutionary memory of structural plasticity that persists even when the physical coordinate is constrained by the crystal lattice. Consistent with this view, crystallographic alternate conformations selected by different lattice environments have been shown to coincide with the conformations intrinsically sampled in solution (Maddipatla et al., 2026). We explicitly test this hypothesis by developing the B-Factor Corrector (BFC), a framework that leverages the deep contextual embeddings of the Evolutionary Scale Modeling 2 (ESM-2) protein language model (Lin et al., 2023). Unlike traditional physical models, BFC treats the recovery of this heterogeneity as a sequence-context-aware translation task that filters out crystal-induced rigidity to reveal the multi-state conformational signal underlying the raw measurement. The conformational diversity captured by structural ensembles in databases such as PDBFlex (Hrabe et al., 2016) reflects spatial variance across crystallographic conditions rather than temporal dynamics from simulations, leaving open whether patterns learned from such data generalize to solution-state dynamics. Our results indicate that static crystallographic data, when interpreted with a language model, contains latent signals governing protein motion. We show that these sequence-decoded signals generalize beyond the crystallographic training distribution to an independent library of molecular-dynamics trajectories, where they mark the residues that switch and are displaced between conformational states; that they spatially localize the conformational transitions underlying disease-associated and chaperone function that evade both crystallography and equilibrium simulation; and that the model organizes them into an interpretable dictionary of heterogeneity features. Building on this, we further establish BFC as an actionable physical prior for computational structural biology, providing a flexibility prior for docking that is statistically equivalent to experimental B-factors and enabling the differential-restraint refinement of predicted protein interfaces.

## Results

### Sequence context enables robust reconstruction of ensemble-RMSF profiles

We first investigated whether a language model could learn the mapping between noisy crystallographic data and solution-state behavior. We benchmarked BFC on a dataset of proteins containing both crystal structures and corresponding ground-truth conformational-heterogeneity profiles from PDBFlex (Hrabe et al., 2016), where per-residue ensemble RMSF is computed from the coordinate variance across multiple depositions of the same protein (see Methods). Comparison with these ensemble-based ground truths highlighted discrepancies in the crystallographic data. Normalized experimental B-factors exhibited low correlation with the ensemble RMSF (Pearson Correlation Coefficient, PCC = 0.15) in the test set (Figure 1B). This suggests that experimental B-factors, often influenced by lattice contacts, may diverge from intrinsic flexibility.

**Figure 1:**
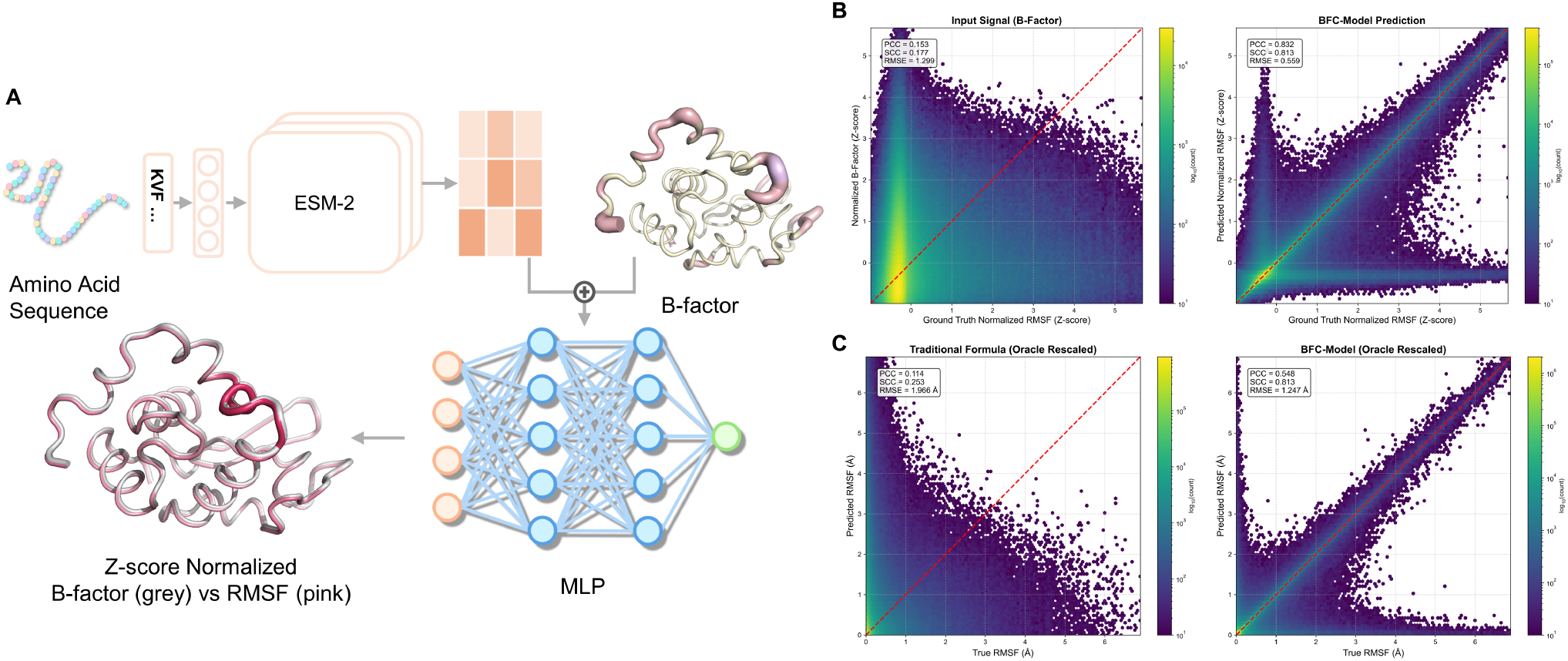
Sequence-context-aware decoding enables quantitative recovery of ensemble RMSF from crystallographic B-factors. **A**, Schematic of the B-Factor Corrector (BFC) workflow. The model extracts deep sequence embeddings via a fine-tuned ESM-2 and combines them with experimental B-factors to predict intrinsic conformational heterogeneity. The structural visualizations illustrate the data transformation. The top structure displays the raw input B-factor distribution, while the bottom superposition reveals the discrepancy between the normalized input B-factors (gray tube) and the groundtruth ensemble RMSF (pink gradient). **B, C**, Global performance evaluation on the test set. Density scatter plots compare the correlation of predictions against ground-truth ensemble RMSF values. **B**, In normalized space (Z-score), BFC (right) denoises the input B-factor signal (left), recovering a high correlation (PCC=0.83). **C**, Evaluation of profile reconstruction in physical units (Å ). To isolate profile fidelity from experimental amplitude variance, both the Traditional Formula (left) and BFC (right) were rescaled to match the ground-truth moments (Oracle Rescaling). Even with correct global scaling, the traditional formula fails to capture the relative fluctuation ranking (SCC=0.25), whereas BFC accurately reconstructs the physical landscape (SCC=0.81), establishing a linear correlation with solution-state reality absent in raw data.

In contrast, BFC recovered a signal consistent with the consensus conformational heterogeneity of the structural ensembles, yielding a PCC of 0.83 on the test set (Figure 1B). We employed an oracle rescaling strategy to isolate profile fidelity from experimental amplitude variance and assess the recovery of physical magnitudes. This protocol standardizes both baseline and model predictions to match the first and second moments of the ground-truth ensemble and thereby focuses the evaluation on the shape of the fluctuation landscape. Even with this global scaling correction, the traditional physical formula failed to reconstruct the relative distribution (Spearman Correlation Coefficient, SCC = 0.25; PCC = 0.11), indicating that the B-factor pattern remains fundamentally distorted. Conversely, BFC achieved a robust reconstruction of the residue-wise conformational-heterogeneity ranking (SCC = 0.81) despite the heavy-tailed distribution of extreme physical fluctuations that dampens linear correlation (PCC = 0.55), and yielded a lower Mean Absolute Error (0.11 Å vs. 0.56 Å, Figure 2B); extended per-protein reconstructions spanning a range of chain lengths are shown in Figure S1. These results suggest that BFC decodes the intrinsic dynamic sequence pattern latent within the static crystal structure, correcting local distortions that analytical formulas cannot address.

**Figure 2:**
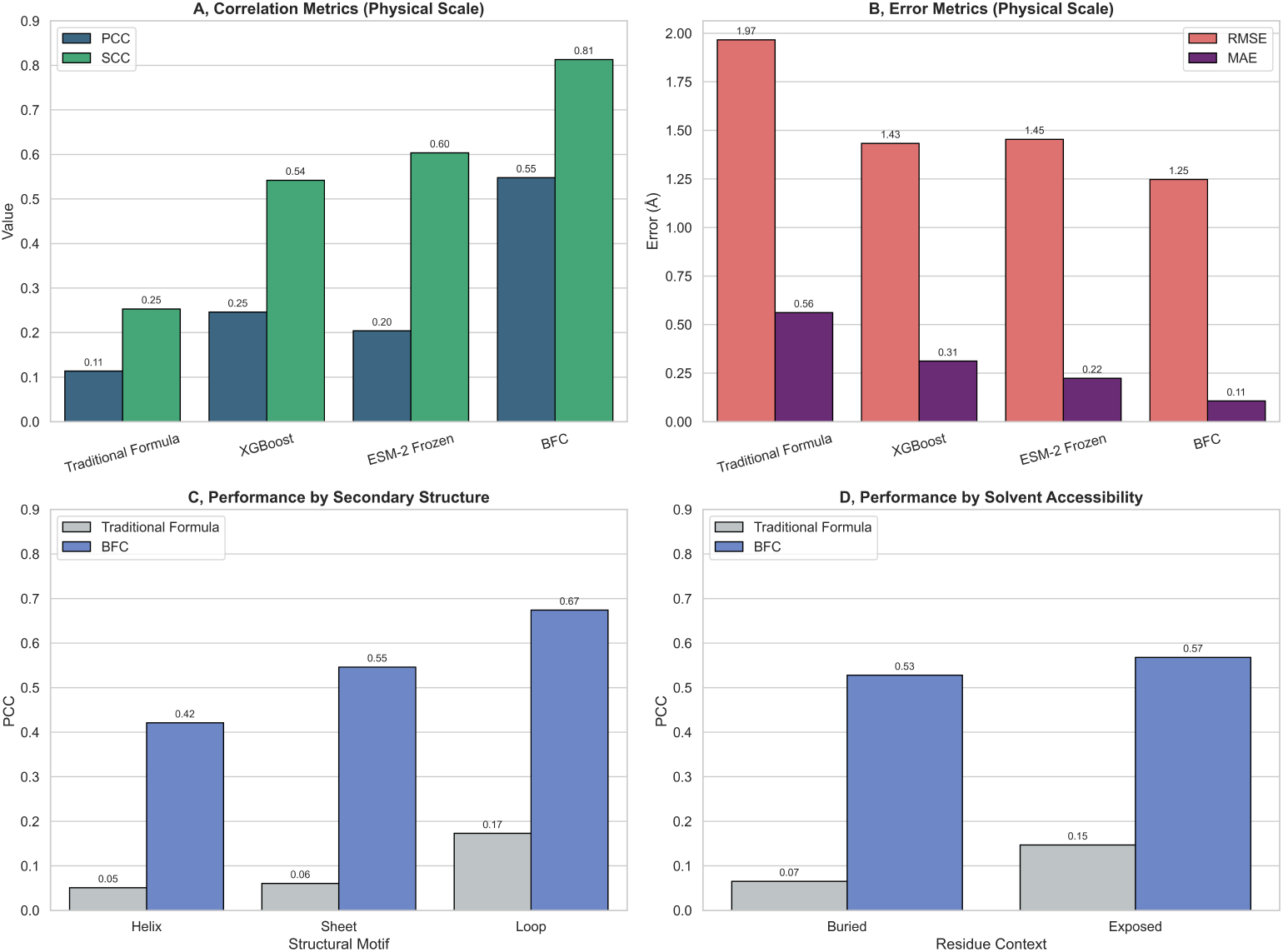
BFC outperforms baselines and demonstrates robust generalization. **A, B**, Systematic benchmarking of dynamic prediction accuracy. BFC (fine-tuned) outperforms the Traditional Formula, XGBoost, and pre-trained ESM-2 (frozen) in both correlation metrics (PCC/SCC) and error metrics (RMSE/MAE). The performance gap underscores the value of end-to-end task adaptation. **C, D**, Performance stratified by structural context. BFC maintains robust performance across secondary structures (C) and solvent accessibility levels (D), achieving high accuracy even in flexible loops and solvent-exposed regions where traditional physical formulas typically fail.

### End-to-end fine-tuning captures biophysical nuances

To understand the biophysical basis of this performance, we isolated the contribution of sequence context by comparing BFC against baseline models (Figure 2A,B; per-method scatter against the ensemble ground truth in Figure S2). A gradient-boosted decision tree (XGBoost (Chen and Guestrin, 2016)) trained solely on B-factor values did not improve correlation, implying that the relationship between B-factors and RMSF is likely not a simple non-linear mapping and that experimental data alone, without sequence context, may be insufficient for denoising. We also compared BFC (fully fine-tuned) with a frozen ESM-2 model, where only the output head was trained. The frozen model yielded limited improvements, suggesting that general-purpose evolutionary representations may not be inherently aligned with specific physical dynamics. To directly quantify the contribution of the experimental B-factor channel, we further compared the full BFC model against a sequence-only ablation trained under identical conditions (Table S1). Removing the B-factor channel consistently degraded performance across all metrics, yielding a small but highly significant net-positive effect for the full model (one-sided Wilcoxon signed-rank test, *p* = 6.3 × 10^*−*13^); this degradation is amplified among the most conformationally heterogeneous residues. Stratifying all test-set residues by their ensemble RMSF (Table S2), we find that the benefit of the experimental channel concentrates toward the most flexible positions. This trend is clearest in the per-residue error metrics, which are unaffected by the range restriction that suppresses correlation within narrow strata. Relative to the sequence-only ablation, the error reduction more than doubles from all residues to the top 0.5% (MAE, +1.9% to +5.2%; RMSE, +2.5% to +3.2%). The correlation gains increase monotonically over the same range, reaching +14.4% (PCC) and +10.0% (SCC). This confirms that the experimental channel contributes orthogonal information beyond the sequence prior, precisely where conformational states are governed by the physical environment rather than by evolutionary sequence conservation.

The performance gap achieved by end-to-end fine-tuning suggests that the model adapts its attention mechanisms to encode sequence-dependent physical properties. To confirm that this performance stems from genuine residue-wise precision rather than global scaling artifacts, we stratified the evaluation by local structural context. This stratification shows that BFC learns distinct dynamic signatures at the residue level. Although buried residues exhibit slightly lower baseline fluctuations (mean Z-score −0.05) compared to solvent-exposed regions (mean Z-score 0.06), the model maintains high predictive accuracy within both regimes. It retains reasonable accuracy in buried hydrophobic cores (PCC = 0.53 vs. 0.07 for the traditional formula, Figure 2D), likely by recognizing the stability conferred by high packing density. Conversely, in solvent-exposed loops, typically enriched with flexible residues such as glycine and proline (Chou and Fasman, 1974; Krieger et al., 2005), the model shows its largest gain over the traditional formula (PCC = 0.67 vs. 0.17, Figure 2C).

Residue-level diagnostic analysis clarifies the physical logic of the model (Figure S3). Although qualitative prediction-ground truth disagreements (residual |*r*| *>* 2.0) affect only 1.64% of all residues, a detailed examination of these outliers resolves a distinction between local intrinsic flexibility and global conformational heterogeneity. When BFC predicts rigidity despite a high ground-truth ensemble RMSF, analysis reveals that these residues typically exhibit low experimental B-factors within their individual crystal structures. This indicates that the high ensemble variance originates from rigid-body domain motions across different crystal forms rather than intrinsic local unfolding. BFC correctly resists being misled by these global spatial translations, accurately decoding the sequence-encoded local stability. Conversely, positive outliers where BFC predicts high flexibility contrary to rigid experimental signals reveal cases of crystallization bias, where BFC identifies latent intrinsic flexibility that happens to be uniformly constrained across available crystal structures.

### Denoising crystal packing artifacts in antibody CDR loops

This decoding approach is illustrated by a test-set case in which it flags and corrects a misleading structural signal. In the crystal structure of an antibody fragment (PDB: 3efd (Uysal et al., 2009)), crystal packing interactions artificially stabilize the Complementarity-Determining Region (CDR) loops, leading to low B-factors that falsely suggest rigidity (Spoendlin et al., 2025; Fernández-Quintero et al., 2021). A naive interpretation of the B-factors would fail to capture the functional plasticity required for antigen binding (Blackler et al., 2022; Fernández-Quintero et al., 2019; Liu et al., 2024).

Guided by the sequence context of the CDR loops, BFC identified the experimental rigidity as a potential artifact. The model predicted a high-flexibility profile that aligned with the ground-truth ensemble variability, correcting the discrepancy in the input signal (Figure 3). This correction suggests that BFC does not merely smooth the input data but captures sequence-dependent physical properties. The model appears to weigh sequence-encoded intrinsic propensities against experimentally observed lattice constraints, helping to recover solution-state dynamics masked by the crystallization process.

**Figure 3:**
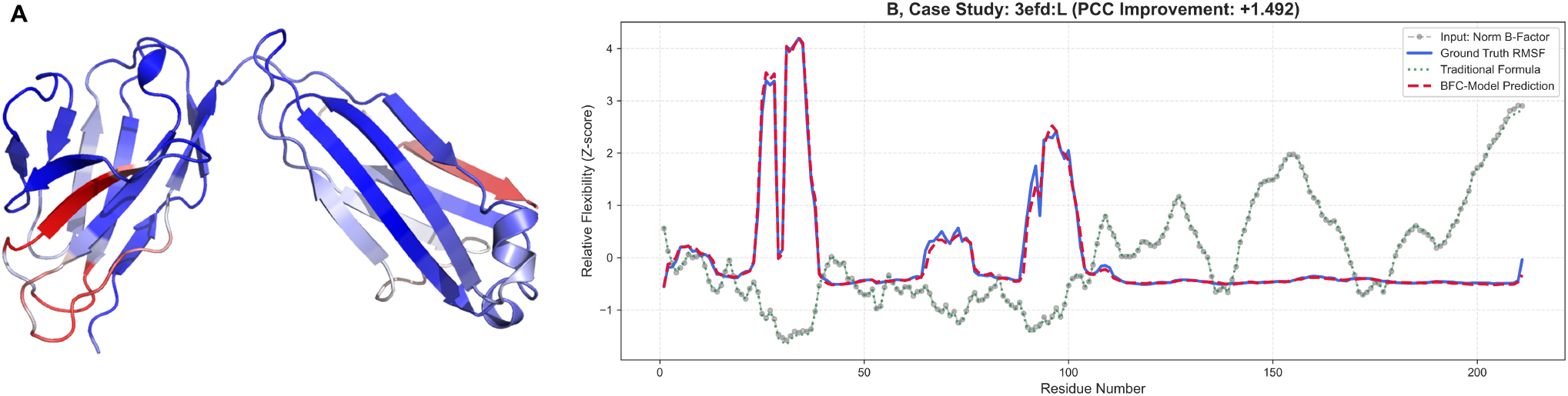
Correction of crystal packing artifacts in antibody CDR loops (PDB: 3efd). Comparison of normalized flexibility profiles (B) reveals a major discrepancy in the CDR loop regions. Experimental B-factors (gray connected dots) falsely suggest rigidity due to crystal contacts, and the Traditional Formula (green dotted line) inherits the same distortion as a monotonic transform of the B-factors. BFC (red dashed line) uses sequence context to identify these artifacts, recovering the high-flexibility profile that matches the ground-truth ensemble RMSF (blue solid line). The structural map (A) visualizes the predicted flexible regions (red) versus rigid scaffolds (blue).

### BFC peaks localize to multi-state conformational regions in an independent MD benchmark

The preceding results establish that BFC recovers ensemble heterogeneity from crystallographic data, but the training signal, PDBFlex ensemble RMSF, reflects conformational diversity sampled across different crystal forms rather than time-resolved motion along a trajectory. It therefore remains to be shown that the regularities BFC extracts from static structural ensembles transfer to explicit-solvent molecular dynamics. To test this, we evaluated BFC on the full set of 1,938 proteins from the ATLAS molecular dynamics simulation database (Vander Meersche et al., 2024) of all-atom explicit-solvent MD simulations. The reference quantities here, per-residue properties of the MD trajectories, constitute a different class of physical observable from the cross-crystal-form ensemble RMSF that BFC was trained on, and were never seen during training, providing an orthogonal ground truth. For each protein we compared the BFC profile predicted from its single crystal structure against per-residue quantities computed from the MD trajectories, testing whether it correctly localizes the residues that undergo conformational rearrangement.

Across the benchmark, BFC does not reproduce the residue-wise MD amplitude; its profile correlates only weakly with trajectory RMSF, with the effective number of sampled conformational states (N_eq_, a local backbone structural-alphabet diversity), and with backbone dihedral entropy (S_*φψ*_, the spread of the per-residue *φ/ψ* distribution), with per-protein median Spearman between 0.15 and 0.21 across channels (Figure 4A,B). This is the expected signature of a positional rather than an amplitude predictor. BFC is dimensionally distinct from any single MD observable, so a low global correlation is expected rather than a failure. What transfers instead is *position*. BFC peaks co-localize with the high regions of every geometric MD channel (RMSF, N_eq_, and S_*φψ*_) over a within-chain permutation null, significant at every threshold, whereas the identical test on shuffled BFC profiles is null throughout (Figure 4C,D; region-overlap enrichment of 1.9– 3.3× at the top 5%, all *p <* 2 × 10^*−*4^). The same enrichment holds for a fully independent measure, AlphaFold predicted Local Distance Difference Test (pLDDT) (Figure 4C,D), indicating that the regions BFC flags are recognizable both from MD geometry and from an orthogonal sequence-based predictor. Ranking the channels by co-localization strength reveals a clear dissociation (Figure 4F). BFC peaks co-localize with every geometric/conformational channel (RMSF, N_eq_, S_*φψ*_, pLDDT), but do not track per-residue interaction-energy fluctuation at all (*σ*_*E*_, CV_*E*_; ∼ 1.0×, indistinguishable from null). BFC thus marks where a backbone samples multiple conformational states, not where local energetics are unstable. Such multi-state behavior is prevalent here. A free-energy landscape analysis finds that 81% of the proteins populate two or more basins (Figure 4H), so the conformational heterogeneity BFC is trained to predict is abundantly present in the independent MD data.

**Figure 4:**
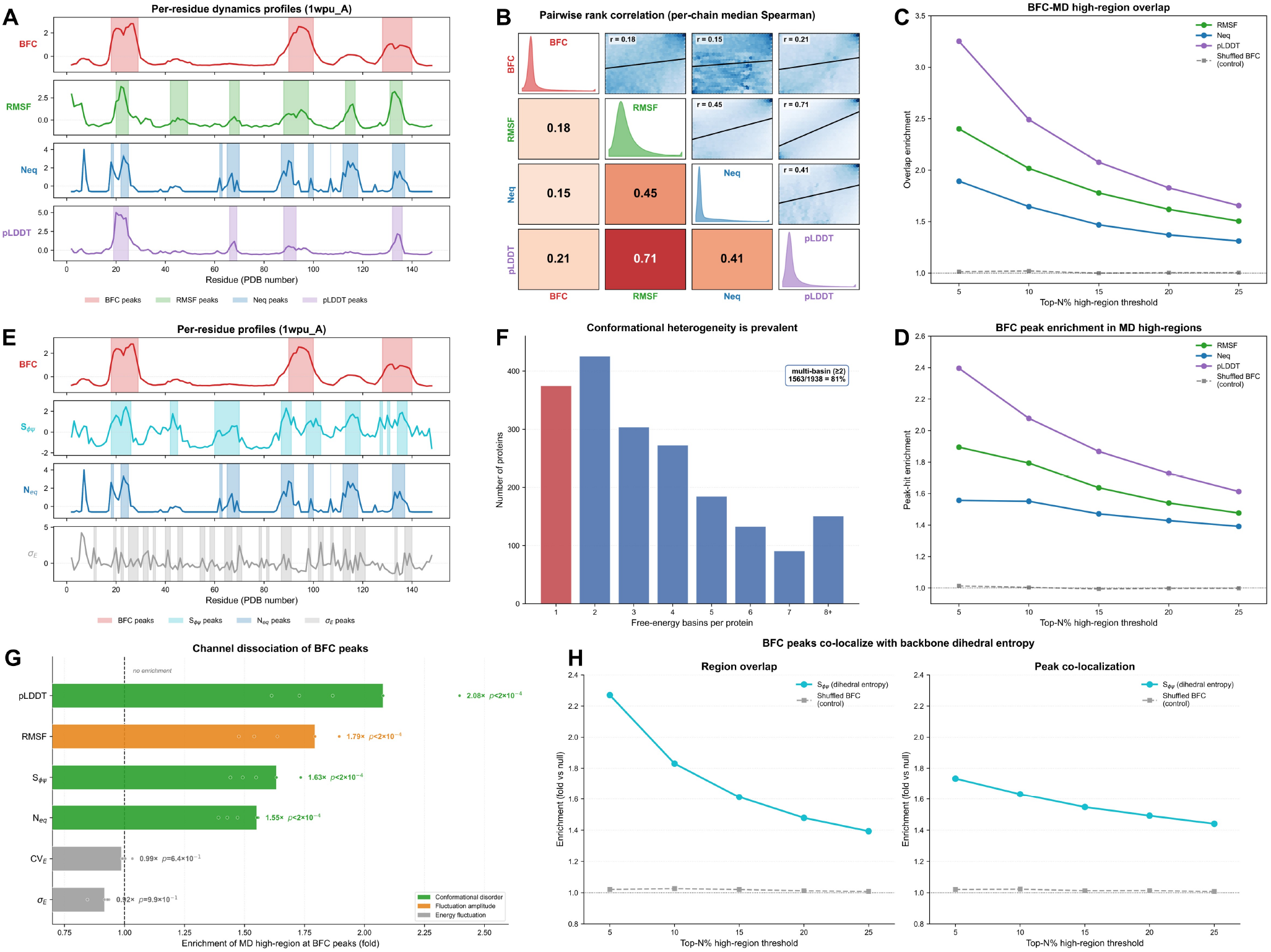
BFC peaks co-localize with conformational-disorder hotspots across an independent MD benchmark (1,938 proteins). **A**, Representative per-residue profiles (1WPU A, HutP antitermination protein) of BFC and three MD-derived quantities (trajectory RMSF, the conformational-state count N_eq_, and AlphaFold pLDDT, a sequence-derived confidence score), each z-scored; shaded bands mark the peaks of each track. **B**, Pairwise rank-correlation matrix (per-protein median Spearman): BFC correlates only weakly with every MD quantity (0.15–0.21), far below their mutual correlations, so BFC is dimensionally distinct from any single MD observable. **C**, Region-overlap enrichment (symmetric Jaccard) between BFC high regions and MD high regions across top-*N* % thresholds, for RMSF/N_eq_/pLDDT versus a shuffled-BFC control; enrichment is significant at every threshold (*p <* 2 × 10^*−*4^, within-chain permutation). **D**, Enrichment of MD high-region residues at BFC peaks (peak co-localization) across the same channels and thresholds. **E**, Representative profiles (1WPU A) of BFC with backbone dihedral entropy S_*φψ*_, N_eq_, and per-residue interaction-energy fluctuation *σ*_*E*_; BFC/S_*φψ*_/N_eq_ peaks coincide while *σ*_*E*_ peaks do not. **F**, Channel dissociation: BFC-peak co-localization (top-5% region-overlap enrichment) across six MD channels. All geometric/conformational channels (RMSF, N_eq_, S_*φψ*_, pLDDT) are enriched (1.9–3.3×), whereas interaction-energy fluctuation (*σ*_*E*_, CV_*E*_) is not (∼ 1.0×, n.s.). BFC marks conformational multi-state character, not energetic instability. Points show individual thresholds; the dashed line marks no enrichment. **G**, The same co-localization holds for S_*φψ*_ (region overlap, left; peak co-localization, right), which we compute directly from the trajectories; because *N*_eq_ and S_*φψ*_ both measure local backbone dihedral diversity (per-protein median Spearman 0.62), this is a robustness check through an independent pipeline rather than a statistically independent replication. **H**, Distribution of free-energy basin counts per protein (single-basin, red; multi-basin, blue); 81% populate ≥ 2 basins. All co-localization tests use a within-chain permutation null; exact effect sizes and *p*-values for all channels and thresholds are tabulated in Table S3.

Having established *where* BFC peaks fall, we asked whether those positions are dynamically active over the simulation timeline, information that the time-averaged disorder measures (*N*_eq_, S_*φψ*_) discard. Assigning each residue’s backbone to Ramachandran basins (*α*/*β*/*α*_*L*_) within each replica and counting confirmed basin transitions, we find that the fraction of residues that switch backbone state over the trajectory rises monotonically with BFC score across all deciles (Figure 5A,B; from 15.9% in the lowest decile to 30.4% in the highest, a 1.91× enrichment; within-chain permutation *p <* 2 × 10^*−*4^). Because this is a transition count over a fixed 100 ns window, we read it as a relative propensity for state switching rather than an absolute kinetic rate, the appropriate reading for a quantity inferred from a single static structure. We next decomposed each residue’s total MD fluctuation into an intra-state component (thermal motion within one basin) and an inter-state component (displacement between global conformational states), on the 592 proteins with clearly separated deep basins (inter-basin barrier ≥ 2.5 kT; the decomposition is ill-posed elsewhere, where the two components are collinear). BFC peaks significantly enrich the residues that bear conformational displacement: both components are enriched at BFC peaks by ∼ 2× over a within-chain permutation null (top 5%: interstate 2.14×, intra-state 2.41×; *p <* 2 × 10^*−*4^ at every threshold; Figure 5C,D). BFC does not separate the two, consistent with its training target, ensemble RMSF, being an aggregate of all cross-conformer displacement rather than a decomposition into thermal versus cross-state motion. Finally, we tested whether this transfers to functionally driven conformational change using an independent set of 118 apo–holo protein pairs (curated from CryptoSite (Cimermancic et al., 2016) and PocketMiner (Meller et al., 2023)), for which the per-residue conformational change upon ligand binding is quantified by comparing the apo and holo crystal structures (1−local CAD-score (Olechnovič et al., 2013)). From the apo structure alone, which BFC has never seen in its bound state, BFC peaks co-localize with the residues that actually change conformation on ligand binding (3.3× region-overlap and 2.7× peak enrichment at the top 5%, *p <* 2 × 10^*−*4^; Figure 5E). A representative case, the small GTPase ARF1, illustrates this directly. The residues BFC scores highest on the apo structure coincide with the switch region that reorganizes between the apo and holo states (Figure 5F). Together, these analyses indicate on independent data that BFC generalizes from static structural ensembles to explicit-solvent dynamics and to ligand-induced conformational change, by enriching for the residues that switch between, and are displaced across, conformational states.

**Figure 5:**
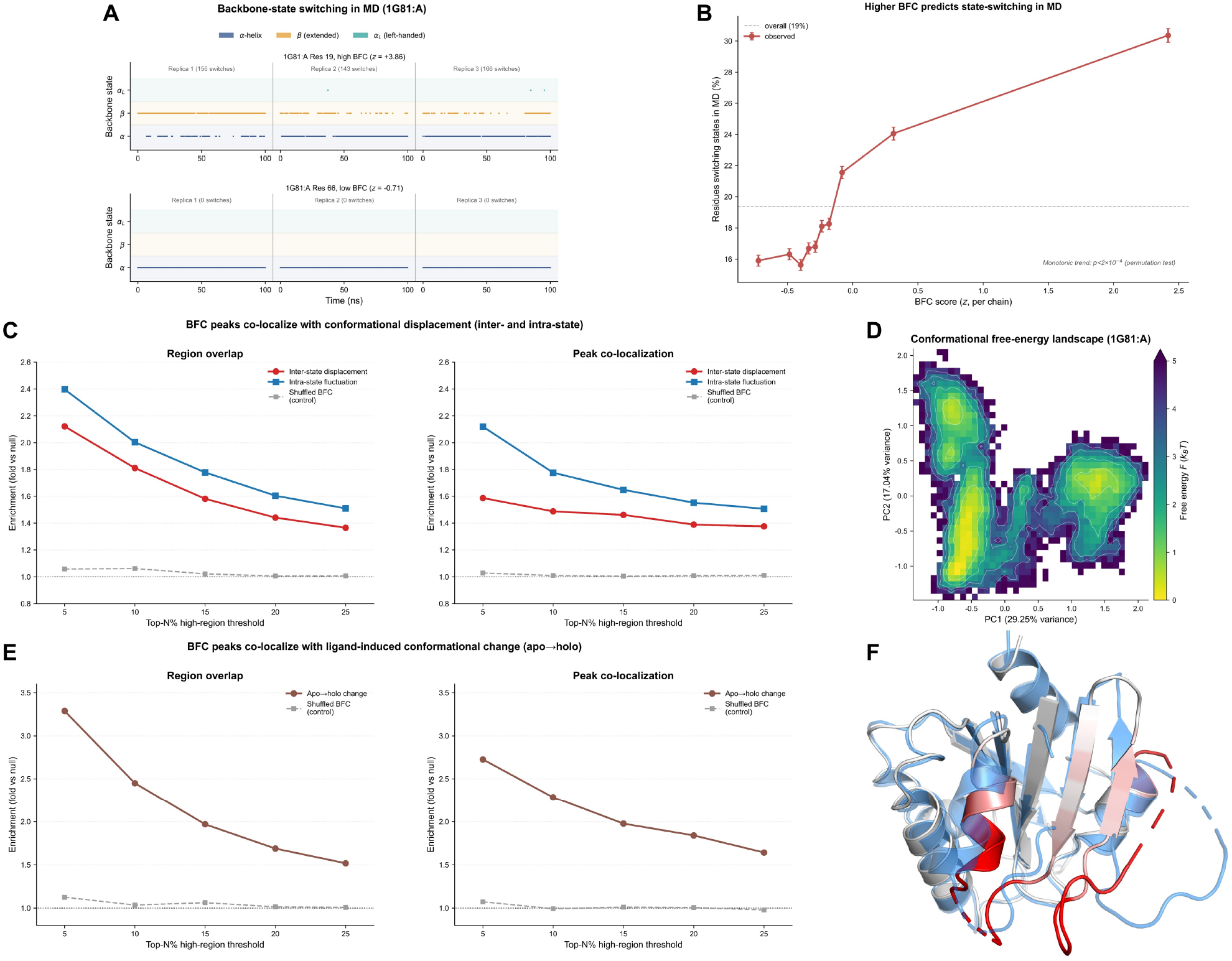
BFC peaks mark residues that switch and are displaced between conformational states, and that reorganize upon ligand binding. **A**, Backbone-state occupancy over time for two residues of one protein (1G81 A, chorismate lyase), across three independent replicas: a high-BFC residue (19; *z* = +3.86) repeatedly switches between *α* and *β* states, whereas a low-BFC residue (66; *z* = −0.71) stays locked in a single state. Illustrative example; the statistical claim is panel B. **B**, Probability calibration: residues binned by BFC score (equal-frequency deciles) versus the fraction that undergoes conformational-state switching in MD. Switching probability rises monotonically with BFC (15.9% in the lowest decile to 30.4% in the highest, a 1.91× enrichment; within-chain permutation *p <* 2 × 10^*−*4^; termini excluded, 1,930 proteins). BFC predicts switching propensity, not rate. Exact statistics and confidence intervals are tabulated in Table S4. **C**, In the 592 proteins with cleanly separated deep basins (inter-basin barrier ≥ 2.5 kT; the decomposition is ill-posed elsewhere, where inter and intra are collinear), enrichment of BFC peaks at inter-state displacement (red) and intra-state fluctuation (blue) high regions, against a shuffled-BFC null (gray). Both components are enriched by ∼ 2× at the top 5% (inter 2.14×, intra 2.41×; *p <* 2 × 10^*−*4^ at every threshold); BFC does not distinguish them, consistent with its ensemble-RMSF target being an aggregate of all cross-conformer displacement rather than a decomposition into thermal versus cross-state motion. Full enrichment curves are tabulated in Table S5. **D**, PCA free-energy landscape of a representative multi-state protein (1G81 A; *F* = − ln *P* over the first two principal components), showing multiple well-separated basins, the conformational distribution sampled in MD. **E**, Generalization to ligand-induced conformational change: across 118 apo–holo pairs (CryptoSite/PocketMiner), enrichment of BFC peaks (predicted from the apo structure alone) at the residues that actually change conformation on ligand binding (1−local CAD-score), against a shuffled-BFC null. Region overlap 3.3× and peak co-localization 2.7× at the top 5% (*p <* 2 × 10^*−*4^); full curves in Table S6. **F**, Representative case: the small GTPase ARF1 (apo, PDB 1RRG), cartoon coloured by BFC score (white, low; red, high), with the ligand-bound holo state (PDB 1S9D) overlaid (blue, semi-transparent). BFC-high (red) regions coincide with the switch region that reorganizes between the apo and holo states.

A natural question is how BFC relates to recent deeplearning ensemble generators that sample multiple conformers from a single structure (e.g. BioEmu (Lewis et al., 2025), ESMFlow/AlphaFlow (Jing et al., 2024)). Across 300 held-out proteins—87 in the intersection of our test set and ATLAS, plus 213 sampled at random from ATLAS (Figure S4)—the observables are only partially coupled. Generated-ensemble per-residue fluctuation aligned preferentially with trajectory-derived MD-RMSF across all 300 proteins (per-protein median Spearman 0.73–0.77), whereas on the 87 proteins that additionally carry PDBFlex ground truth its agreement with the cross-deposition structural heterogeneity that BFC targets was far weaker and comparable to the MD-vs-heterogeneity baseline itself (0.20–0.26 against PDBFlex, versus ATLAS-MD-vs-PDBFlex 0.21); BFC showed the opposite profile (0.16 with MD-RMSF, 0.60 with PDBFlex ensemble RMSF). This dissociation reflects accessible information rather than model capacity. A single input structure, whether resampled by a generator or reduced to elastic-network geometry (ANM), does not preferentially recover the between-deposition displacement that defines cross-crystal heterogeneity, which BFC reaches through evolutionary sequence context. These generators emulate the accessible molecular-dynamics equilibrium ensemble, and even when used to seed molecular simulations they do not reproduce the experimentally observed sidechain conformational heterogeneity that governs transitions in systems such as glycine transporter 1 and plasmepsin-II (Bhakat and Strauch, 2026). Among residues that are heterogeneous across depositions yet static within any single crystal (buried and secondary-structured; Figure S4), BFC assigns high scores while B-factor and ANM do not (per-protein paired Wilcoxon, BFC *p* = 4.7 × 10^*−*12^ positive; B-factor *p* = 5.3 × 10^*−*9^ and ANM *p* = 7.2 × 10^*−*16^ reversed). BFC and ensemble generators are therefore complementary rather than competing. A generator proposes 3D conformers along the thermally accessible landscape, whereas BFC supplies a per-residue amplitude of cross-deposition heterogeneity and does not itself generate structures.

### Deciphering disease-critical dynamics latent in the PDB

To validate the utility of our model for biological discovery, we applied BFC to the ATLAS database (Vander Meersche et al., 2024), an external repository of molecular dynamics simulations. This dataset is distinct from our ensemble-based training distribution, allowing us to test whether the model generalizes from static structural clusters to time-resolved dynamic trajectories. We analyzed the SH2 domain of SLAM-associated protein (PDB: 1D4T (Poy et al., 1999)), a critical signaling module. Consistent with the functional nature of this region, the experimental B-factors for the loop spanning residues 68–73 exhibited a distinct peak of flexibility (Figure 6C).

**Figure 6:**
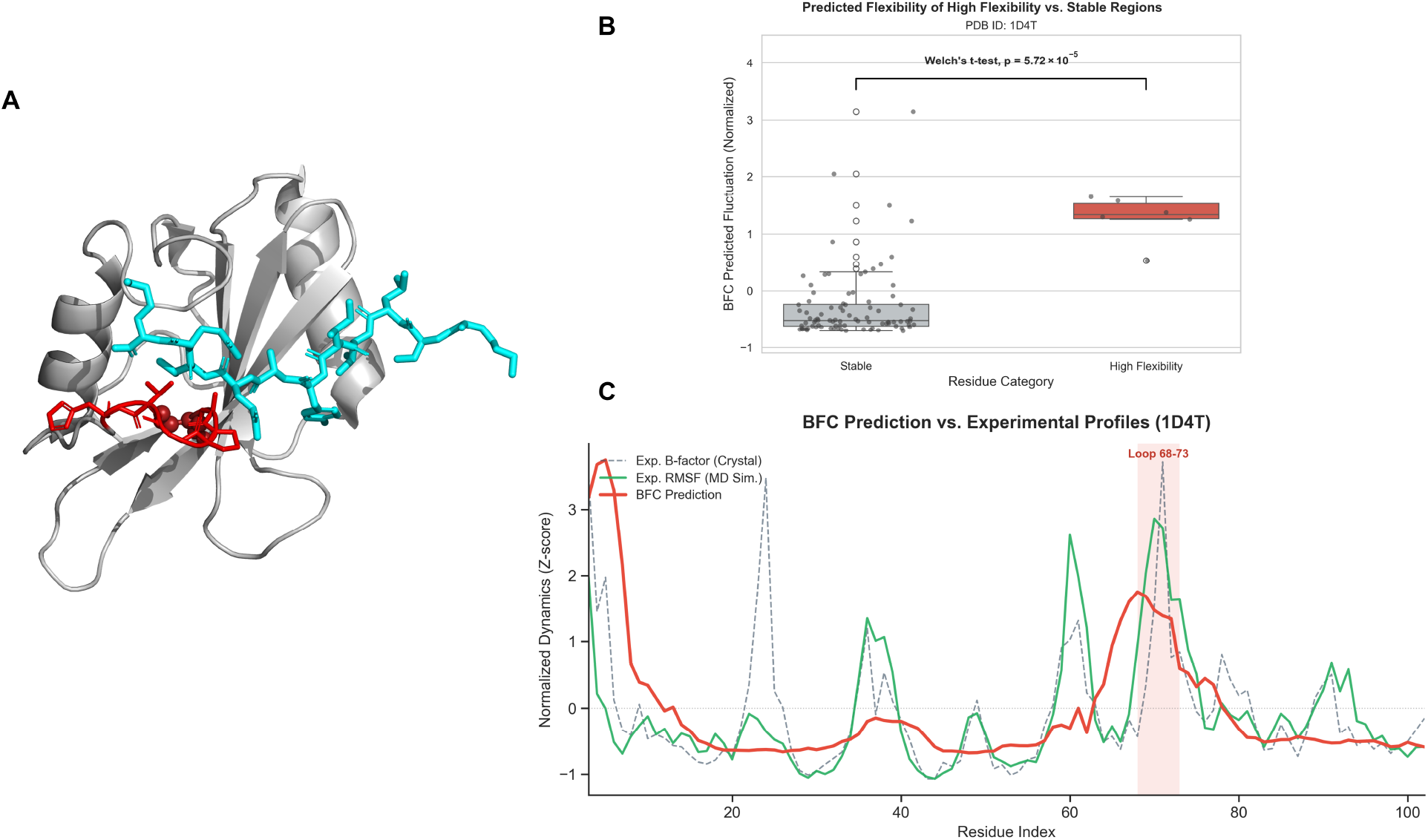
Identification of disease-associated flexible motifs (PDB: 1D4T). **A**, Structural visualization of the SH2 domain where BFC identifies a highly flexible loop (residues 68-73, red sticks) accommodating the SLAMF1 ligand (cyan). The Arg68 residue (red spheres) associated with XLP1 disease resides within this flexible region. **B**, Statistical validation indicating that BFC-predicted flexibility for the binding loop is significantly elevated compared to the stable regions (Welch’s *t*-test, *p* = 5.72 × 10^*−*5^). **C**, Comparative flexibility profiling against external molecular dynamics simulations. The BFC prediction (red solid line) is overlaid with experimental crystallographic B-factors (gray dashed line) and ground-truth RMSF derived from the ATLAS MD database (green solid line). BFC accurately recovers the solution-state flexibility peak at the binding loop (residues 68-73) consistent with the MD trajectory, confirming the capacity of the model to generalize from structural ensembles to time-resolved simulations and pinpoint pathogenicity-relevant dynamics.

BFC accurately reproduced this localized peak of flexibility. Structural analysis indicates that this loop acts as a gatekeeper for the binding of the SLAMF1 ligand (Figure 6A) (Poy et al., 1999). The Arg68 residue, located at the apex of this predicted flexible region, is the site of a missense mutation associated with X-linked lymphoproliferative syndrome type 1 (XLP1) (Booth et al., 2011). Statistical analysis confirmed that the BFC-predicted flexibility for this loop was significantly higher than the scaffold background (Welch’s *t*-test, *p* = 5.72 × 10^*−*5^), identifying a functional hotspot and indicating that the sequence-contextualized flexibility signal generalizes to disease-relevant solution-state dynamics (Figure 6B,C). This capacity to pinpoint function-critical dynamics suggests that BFC can help re-analyze existing PDB entries to generate hypotheses about protein function and disease mechanisms. Two additional examples following the same ATLAS-validated approach, azurin (PDB: 1CUO) and barstar (PDB: 1B2S), likewise resolve functional-loop flexibility (Figure S6 and Figure S7).

### Unmasking cryptic conformational flexibility latent in static structures

While the previous case demonstrated consistency between BFC and molecular dynamics, a more profound application lies in identifying functionally relevant flexibility that evades detection by crystallography and standard equilibrium simulations. We analyzed the Rubisco accumulation factor 1 (RAF1, PDB: 4WT3), a critical chaperone for plant photosynthesis (Hauser et al., 2015). Standard structural metrics present a limited view in which both crystallographic B-factors and ATLAS MD simulations depict the *α*-domain as a rigid scaffold (Fig. 7A). BFC conversely predicts a dominant high-intensity flexibility peak localized to the C-terminal cap of Helix *α*2 and its adjoining loop, apexing at a conserved glycine (G112; residues 106–113).

**Figure 7:**
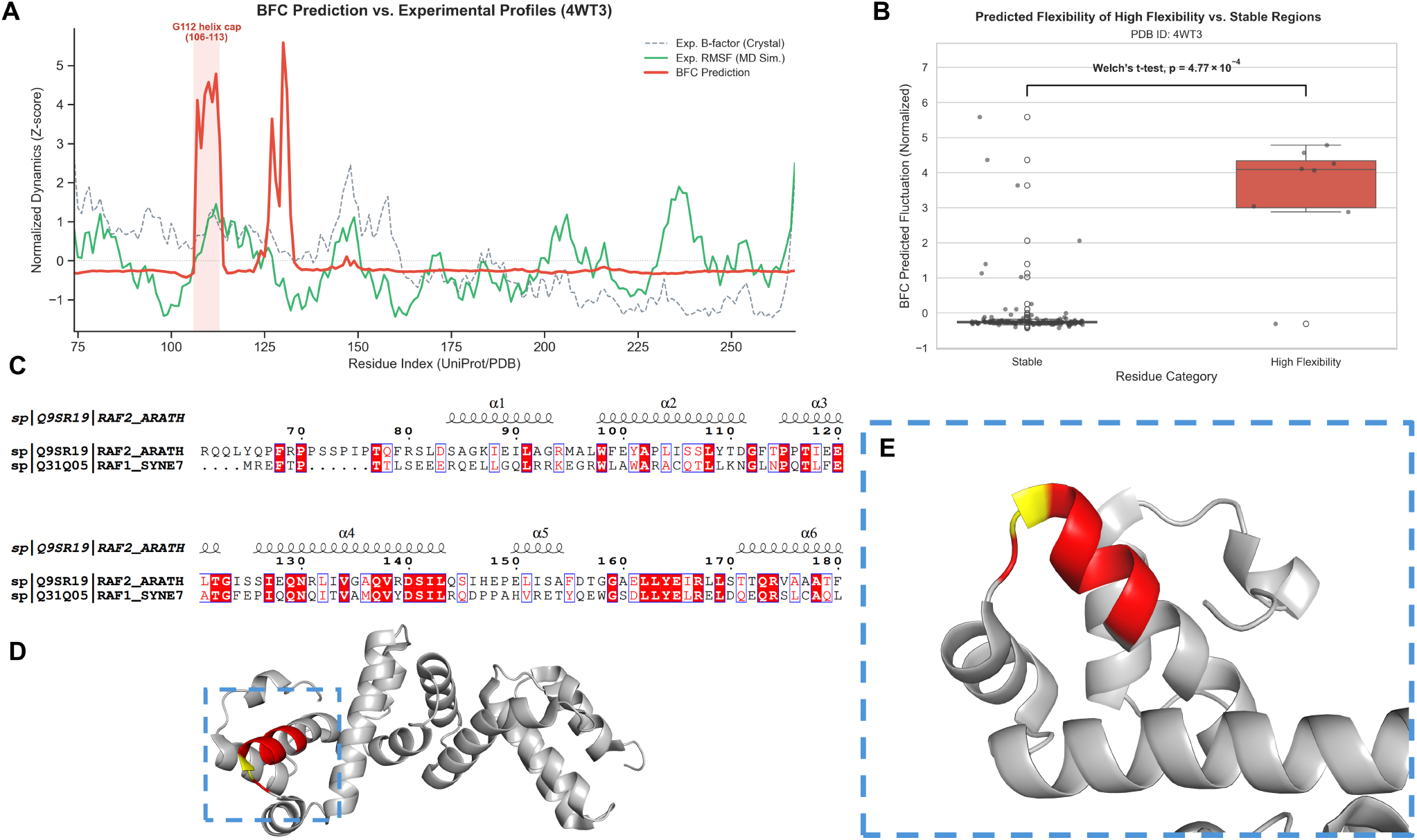
BFC unmasks cryptic conformational flexibility in the Rubisco chaperone RAF1 (PDB: 4WT3). **A**, Comparison of structural metrics where crystallographic B-factors (gray dashed) and standard MD simulations (ATLAS database, green) depict the *α*-domain as uniformly rigid. Conversely, BFC (red) predicts a dominant high-flexibility peak at the C-terminal cap of Helix *α*2, apexing at the conserved glycine G112 (residues 106–113), together with a secondary peak at N130 (within Helix *α*4) analysed in the following section. **B**, Statistical analysis confirming that the predicted fluctuation of this region is significantly higher than the stable background of the protein (Welch’s *t*-test, *p* = 4.77 × 10^*−*4^). **C**, Sequence alignment with the homolog *S. elongatus* Raf1 reveals the conserved nature of the predicted region, where a conserved glycine residue (G112, red box) sits at the terminus of Helix *α*2. **D, E**, Structural mapping of the predicted flexible region. **D**, Overview of the RAF1 *α*-domain monomer. **E**, Zoomed view of the BFC-predicted flexible region (red, residues 106–113) at the C-terminal cap of Helix *α*2. The conserved glycine G112 (yellow sticks) sits at the C-terminal cap of Helix *α*2, where its low *α*-helical propensity and expanded backbone conformational freedom confer a local conformational plasticity that is masked in the static crystal structure yet recovered by BFC from sequence context.

We resolved this discrepancy by performing a sequence alignment between the crystallized *A. thaliana* RAF1 and its functional homolog from *S. elongatus* (Syn7942), which confirms that G112 is strictly conserved. Secondary-structure assignment using the Define Secondary Structure of Proteins (DSSP) algorithm places this glycine in a coil at the C-terminal cap of Helix *α*2, just upstream of Helix *α*3, rather than within a helix body. Glycine has the lowest *α*-helical propensity of the twenty amino acids (Pace and Scholtz, 1998), a preference rooted in the high backbone conformational entropy it retains when, lacking a C*β*, it is not constrained to the helical state (Scott et al., 2007); moreover, a conserved glycine at a helix C-terminus is the hallmark of a canonical helix-termination motif (Aurora et al., 1994). Consistent with the capacity of glycine to introduce local packing defects and backbone flexibility even within a formed helix (Högel et al., 2018), this position carries an intrinsic propensity for local conformational rearrangement at the helix terminus that the crystal lattice suppresses (Fig. 7B,C).

The strict conservation of G112 across RAF1 homologs indicates that this position is under selective constraint. RAF1 acts as an assembly chaperone whose *α*-domain engages the Rubisco large subunit (RbcL) by bracketing each RbcL dimer at its top and bottom edges, with a shallow groove on the conserved surface cradling the back of the RbcL C-terminal domain; this interaction must remain dynamic, because the RAF1 *α*-domain and the small subunit RbcS occupy overlapping sites on RbcL and RAF1 must ultimately be displaced to complete holoenzyme assembly (Hauser et al., 2015). Consistent with their focus on this inter-domain assembly geometry, the structural studies that established this mechanism model the *α*-domain as a rigid body and attribute the requisite dynamics to the disordered *α*–*β* linker (Hauser et al., 2015). The G112 plasticity that BFC decodes is a distinct, intra-domain signal. It is not apparent in either the crystallographic B-factors or the ATLAS MD simulations, yet BFC recovers it from sequence context alone, illustrating the capacity of the model to expose a sequence-encoded, evolutionarily conserved flexibility that static coordinates leave hidden (Fig. 7D,E).

BFC predicts a second peak at N130 that cannot be explained by the local-flexibility mechanism invoked for G112. DSSP places N130 within the body of Helix *α*4, and both MD simulation and cross-structure superposition indicate that this position is locally rigid. This apparent paradox, a high predicted fluctuation at a structurally rigid, conserved residue, motivated a mechanistic interpretability analysis of the model’s internal representations, which we present below.

### BFC learns an interpretable dictionary of conformational heterogeneity features

To understand how BFC arrives at a prediction that contradicts the static coordinates, we examined its internal representations with a sparse autoencoder (SAE) (Huben et al., 2024), which decomposes the dense layer-6 activations of the model into a dictionary of sparse, individually interpretable features. A small subset of these features tracks conformational heterogeneity specifically; their activation correlates with ensemble RMSF while the vast majority of the dictionary does not (Fig. 8A). These features carry a secondary-structure fingerprint, preferentially activating in loops (Fig. 8B), yet their association with heterogeneity is not merely an artifact of secondary structure; it survives stratification within each structural class (Fig. 8D). The sites they mark show conformational heterogeneity at the extreme of the distribution even though their crystallographic B-factors are only moderate (Fig. 8C), mirroring at the feature level the decoupling of intrinsic heterogeneity from the lattice-suppressed crystallographic signal that BFC is trained to correct. The dictionary is further organized along two largely separable axes. One set of features specializes for static structural context (helix, sheet, buried core, or exposed loop), each with a tight amino-acid and solvent-accessibility signature consistent with a monosemantic unit (Fig. 8E,F). Heterogeneity forms a distinct axis that is not simply a byproduct of this structural specialization (Fig. 8E), indicating that conformational heterogeneity is represented within BFC as a signal separable from the static fold.

**Figure 8:**
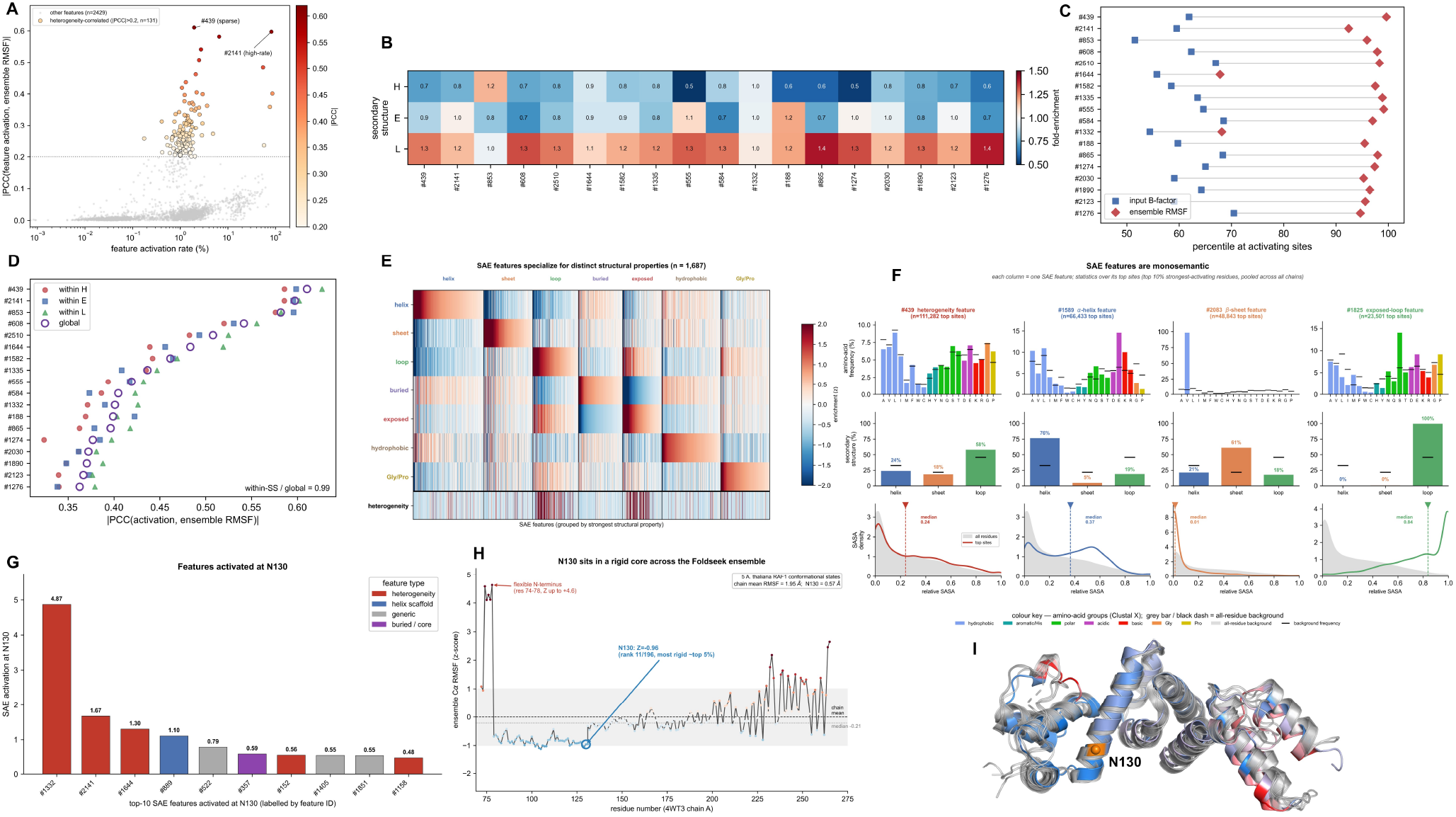
BFC represents conformational heterogeneity through an interpretable, specialized feature dictionary, of which N130 is a representative instance. A sparse autoencoder (SAE; *F* = 2560 features, top-*K* = 64) was trained on BFC layer-6 representations; this configuration was selected from a sweep over dictionary size and sparsity (Figure S5). **A**, Feature atlas: each SAE feature plotted by activation rate (*x*, log scale) against the absolute correlation between its activation and ensemble RMSF, |PCC| (*y*). 131 features exceed |PCC| *>* 0.2 (max 0.610, feature #439), against a background of 2429 features with median |PCC| = 0.011, indicating that heterogeneity is captured by a small, sparse subset rather than diffusely. **B**, Secondary-structure fold-enrichment of the top heterogeneity-correlated features (loop mean 1.23×, helix 0.76×). **C**, Activating sites of these features sit at extreme ensemble-RMSF percentiles (median 97th) but only moderate B-factor percentiles (median 62nd), the gap that BFC is built to recover. **D**, Controlling for secondary structure leaves the feature–heterogeneity correlation essentially unchanged (global |PCC| = 0.449 vs. within-class 0.443; ratio 0.99), excluding a trivial loop-detector confound. **E**, Dictionary-wide specialization: 1687 active features (columns) scored against eight residue properties (rows). The diagonal block structure shows feature groups specialized for distinct structural contexts; the heterogeneity row is an independent axis, enriched in the loop/exposed groups (+0.74/+0.64) and depleted in helix/sheet/buried (−0.16/−0.22/−0.33). **F**, Representative single-feature portraits (top activating sites): #439 (a context-dependent heterogeneity detector), #1589 (*α*-helix, 76%), #2083 (*β*-sheet 61%, Val 98%, mean relative SASA 0.01), and #1825 (exposed loop 100%, SASA 0.84), each showing a tight amino-acid/secondary-structure/SASA signature consistent with a monosemantic unit. **G**, The ten strongest SAE features activated at N130, coloured by annotation: the dominant features are heterogeneity detectors (#1332, #2141, #1644), while a helix-scaffold feature (#889) and a buried-core feature (#357) also activate, reflecting N130’s buried *α*-helical context. **H**, Per-residue ensemble C*α* RMSF (z-score) across an ensemble of five *A. thaliana* RAF1 conformational states (the X-ray query 4WT3:A superposed with four Foldseek structural homologs of ≥95% sequence identity, namely 8IO2:I, 8ILB:E, 8IOJ:E and 8IOJ:F, by iterative rigid-core alignment on the least-variable 60% of residues, which concentrates the superposition on the rigid functional core rather than letting mobile regions dominate the fit (Mukherjee and Hummer, 2026)); N130 lies in the rigid core (RMSF 0.57 Å, *Z* = −0.96, rank 11/196), whereas the flexible N-terminus reaches *Z >* +4. **I**, The five conformers superposed and coloured by ensemble RMSF (marine, rigid; pink, intermediate; red, flexible), with N130 marked (orange sphere) within the rigid core.

This dual structure grounds the N130 case and resolves the paradox raised above. The features that activate most strongly at N130 are the same heterogeneity-tracking features characterized across the dictionary above, whose activation is systematically tied to ensemble RMSF and is not reducible to secondary structure (Fig. 8A,D); their engagement here places N130 within this global population and accounts for its high predicted fluctuation. The same position also activates features whose independent structural annotation marks a buried, *α*-helical context (Fig. 8G). The two readouts are not in conflict because they measure different things. Heterogeneity is judged against ensemble RMSF, whereas rigidity is judged against structural annotation that never enters the model as a label. The structural reading is corroborated geometrically. Across an ensemble of RAF1 conformational states identified by structure search with Foldseek (Van Kempen et al., 2024), N130 occupies the rigid core (RMSF 0.57 Å, *Z* = −0.96, rank 11/196; among the most rigid ∼6% of residues), in contrast to the flexible N-terminus (Fig. 8H,I). N130 is thus best read not as a confirmed high-heterogeneity residue but as a case where sequence context activates the same heterogeneity-tracking features that are library-wide predictors of high ensemble RMSF (Fig. 8A,D), even though the five available RAF1 conformers place the site in the rigid core (*Z* = −0.96; Fig. 8H). Whether the corresponding conformational state is simply unsampled by these few structures, or the sequence-encoded propensity is not realized in this particular protein, cannot be adjudicated from so small an ensemble. What N130 does establish, and what generalizes beyond this single site, is that conformational heterogeneity is encoded within BFC as a signal separable from the static fold (Fig. 8E), rather than as a cherry-picked anomaly.

### Application to molecular docking and interface refinement

Both applications test whether the sequence-denoised BFC signal can serve as an actionable flexibility prior where experimental B-factors are noisy or absent. We assessed whether BFC-predicted flexibility can substitute for experimental B-factors as a residue-level flexibility source across the PDB-bind Refined Set comprising 1,204 complexes (Wang et al., 2005; Liu et al., 2017). When used to nominate flexible pocket residues for docking, crystallographic B-factors risk propagating lattice artifacts; we therefore tested whether the sequence-context-denoised BFC signal yields equivalent docking behavior. Across the 1,204 complexes, the ligand RMSD distribution obtained under BFC guidance was statistically equivalent to that obtained under experimental B-factor guidance (two one-sided tests, *p* = 1.1 × 10^*−*8^ at a 0.5 Å margin; mean difference 0.03 Å ; Fig. 9A). As expected for a redocking benchmark, in which each ligand is returned to its own bound-state pocket, the rigid receptor attained the highest success and added flexibility did not improve accuracy in this setting. A sequence-context-denoised flexibility prior therefore reproduces the behavior of experimental B-factors without introducing artifacts, so BFC can replace them as a flexibility source when the latter are noisy or lattice-dominated.

**Figure 9:**
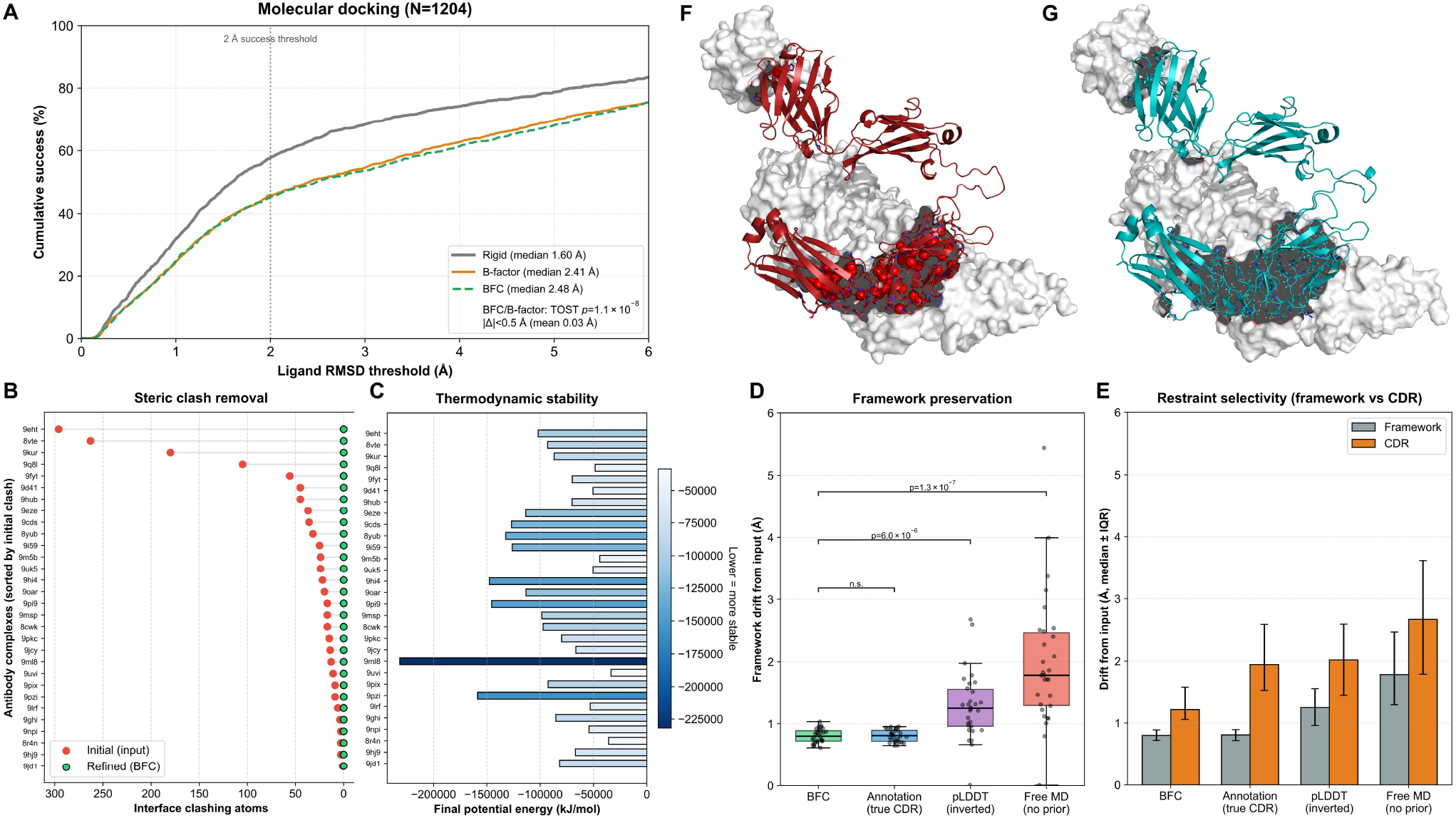
Utility of sequence-derived flexibility for molecular docking and antibody interface refinement. **A**, Molecular docking on the PDBbind refined set (N=1,204). Cumulative success curves (fraction of targets below an RMSD threshold) show that flexibility guided by BFC (green) is statistically equivalent to that guided by experimental crystallographic C*α* B-factors (orange; two one-sided tests *p* = 1.1 × 10^*−*8^ at a 0.5 Å margin, mean difference 0.03 Å ). As expected for a redocking benchmark, where each ligand is returned to its own bound-state pocket, the rigid receptor (gray) attains the highest success, so added side-chain flexibility offers no advantage in this setting; BFC nonetheless reproduces the experimental prior without introducing artifacts. B–G, Proof-of-concept refinement of antibody–antigen interfaces built from ESMFold predictions (30 complexes). **B**,**C**, BFC-guided refinement (BFC scores set differential positional restraints, rigid framework, relaxed CDR loops): **B**, interface clashing atoms per complex before (red) and after (green); **C**, final potential energies, relaxing systems from physically invalid high-energy initial states to stable minima. **D**,**E**, Attribution by four matched restraint schemes on identical MD, comprising BFC-guided, expert-annotation-guided (CDR/framework partition from ANARCI Chothia numbering), unrestrained, and one guided directly by the inverted pLDDT confidence that BFC itself receives as input. **D**, Framework drift from the initial structure is equivalent between BFC and expert annotation (two-sided Wilcoxon *p* = 0.97) and significantly lower than unrestrained MD (*p* = 1.3 × 10^*−*7^), which lets the framework collapse; guiding the same restraints with the inverted pLDDT directly gives a larger framework drift than BFC (median 1.25 vs 0.80 Å, *p* = 6.0 × 10^*−*6^). **E**, Within each scheme, drift of framework versus CDR residues: BFC and annotation selectively immobilize the framework while releasing the CDR loops, whereas unrestrained MD displaces both comparably. **F**,**G**, Structural example (PDB 9EHT). **F**, Initial rigid alignment of the ESMFold prediction (red) to the antigen (white) shows severe steric clashes. **G**, After BFC-guided refinement (teal) the CDR loops adapt to the antigen surface, resolving clashes with high shape complementarity.

As a proof of concept for interfaces that lack experimental B-factors, we asked whether BFC-derived flexibility is an actionable prior for refining antibody-antigen interfaces built from sequence-based predictions. State-of-the-art predictors such as ESMFold (Lin et al., 2023) generate accu-rate but unliganded backbones; rigidly aligning them to an antigen yields severe steric clashes and physically invalid, high-energy interfaces. Because such predictions carry no experimental B-factors, we supplied BFC with per-residue ESMFold pLDDT confidence as a proxy input (see Methods), and used the resulting per-residue scores to set differential positional restraints, immobilizing the predicted-rigid frame-work while releasing the predicted-flexible CDR loops, during molecular dynamics (MD) refinement. Across 30 complexes this eliminated interface steric clashes and relaxed the systems from extremely high initial energies to stable minima (Fig. 9B,C). To attribute the effect to the BFC prior rather than to the MD engine, we compared four matched restraint schemes on identical MD: BFC-guided, expert-annotation-guided (CDR/framework partition from ANARCI Chothia numbering (Dunbar and Deane, 2016)), unrestrained, and one guided directly by the inverted pLDDT confidence that BFC itself receives as input. Framework drift from the initial structure under BFC-guided restraints was statistically indistinguishable from the expert-annotation gold standard (two-sided Wilcoxon *p* = 0.97) and significantly lower than that of unrestrained MD (*p* = 1.3 ×10^*−*7^), which allowed the framework to collapse (Fig. 9D). Guiding the same restraints with the inverted pLDDT directly, rather than with the BFC output, gave a larger framework drift (median 1.25 vs 0.80 Å, twosided Wilcoxon *p* = 6.0 × 10^*−*6^), so the sequence-context correction, not the confidence signal alone, is what stabilizes the framework here. The refinement was selective: BFC and annotation both immobilized the framework while permitting CDR-loop adaptation, whereas unrestrained dynamics displaced both regions comparably (Fig. 9E); the 9EHT complex illustrates the resulting clash-free, shape-complementary interface (Fig. 9F,G). Antibodies are a rare case where an expert flexibility partition (CDR versus framework) exists, providing a gold standard against which a general prior can be validated; that BFC, using no antibody-specific rules, reproduces this expert partition (Wilcoxon *p* = 0.97) is precisely what licenses the same anchor-and-release refinement for the large majority of interfaces that possess no expert annotation. The claim is transferability, not superiority over a domain-specific rule.

## Discussion

Our study indicates that crystallographic B-factors retain usable dynamic signal beneath their environmental and lattice contributions. Treating the B-factor as a convolution of intrinsic motion and environmental constraint rather than as random noise, we use a protein language model as an evo-lutionary prior to decouple the two and to read out conformational heterogeneity directly from sequence context, countering the tendency of structural predictors to memorize static training conformations (Chakravarty et al., 2024). The recovered quantity is not the B-factor itself but a distinct, sequence-encoded heterogeneity profile. A sparse autoencoder resolves it as a dimension separable from static structural context, and the same signal proves integrable downstream, substituting for experimental B-factors in flexible docking and stabilizing antibody frameworks during refinement. This mirrors recent breakthroughs in protein design (Dauparas et al., 2022; Watson et al., 2023; Dauparas et al., 2025; Hayes et al., 2025), where deep learning bridges sequence and physical realizability. We first examine how this signal behaves on independent dynamics, then situate BFC against existing flexibility methods and delineate the scope of these claims.

The signal that BFC decodes from static crystallographic ensembles transfers to independent molecular dynamics simulations, though as a predictor of position rather than of amplitude. BFC does not reproduce the residue-wise magnitude of the trajectory RMSF, and its correlation with any single dynamic quantity derived from the simulations remains low, as expected for observables that differ in physical dimension. The positions it scores highest nonetheless coincide with the residues that are conformationally active in the trajectories, and this agreement holds across several independent facets of the dynamics. The trajectory RMSF, the diversity of sampled backbone states, and the backbone dihedral entropy are all enriched at these positions, as is an orthogonal confidence measure derived from sequence, whereas the per-residue fluctuation of interaction energy is not. This dissociation indicates that BFC locates where a backbone samples multiple conformational states rather than where local energetics are unstable. The same positions are enriched for residues that switch backbone state during a trajectory, for residues that carry displacement between free-energy basins, and, when the prediction is made from the unbound structure alone, for residues that reorganize upon ligand binding. A signal distilled from cross-deposition crystallographic variation therefore locates functional conformational change in data never encountered during training, which suggests that BFC reflects a genuine physical property rather than an artifact specific to crystallization.

BFC recovers conformational heterogeneity at a specific class of residues, namely those that stay buried and secondary-structured within any single form yet vary across crystal depositions, the locally-rigid-but-globally-heterogeneous signature exemplified by N130, which experimental B-factors, elastic-network models, and single-structure ensemble generators alike score as rigid (Figure S4). Such a signal is easily overlooked because conventional structural comparison is dominated by the most mobile loops and disordered termini while the rigid functional core contributes negligibly to a global RMSD (Mukherjee and Hummer, 2026). Deep learning tools that predict flexibility *de novo* from sequence (Ilzhöfer et al., 2022; Hu et al., 2021; Vander Meersche et al., 2025b; Pandey et al., 2023; Yang et al., 2025) or from structure (Song et al., 2024) likewise track the exposed, mobile loops for which the experimental B-factor is already an adequate proxy; the buried heterogeneous core is where sequence context supplies non-redundant information.

BFC reaches this core by taking an experimental B-factor as input and using sequence context to separate its intrinsic conformational-heterogeneity component from crystallographic noise. Such signal-from-noise decomposition of experimental measurements recurs across the life sciences, from cryo-EM density map enhancement (Sanchez-Garcia et al., 2021) to single-cell transcriptome imputation (Eraslan et al., 2019). Because it draws on existing measurements, BFC recovers higher-fidelity dynamics from the more than 190,000 PDB entries that already carry B-factors. Sequence-only structure predictors such as AlphaFold3 instead tend to collapse to a single dominant conformation, and recovering their latent heterogeneity likewise requires conditioning on experimental measurements such as NMR, crystallographic density, or cryo-EM data (Maddipatla et al., 2026).

Several boundaries define the scope of the present work. First, BFC is by construction a corrector rather than a *de novo* predictor, since it requires an experimental B-factor channel as input, and therefore does not apply to structures that carry no such measurement, including purely computational models, unresolved regions, and proteins that resist crystallization such as intrinsically disordered proteins. Second, because BFC draws its corrective signal from the evolutionary representations of a protein language model, its inference is expected to be weakest for sequences that are poor in evolutionary information, such as *de novo* designed proteins, rapidly evolving viral proteins, evolutionarily young proteins, and the hypervariable CDR loops of antibodies, where homology-based priors are known to degrade (Hou et al., 2026); consistent with this boundary, the antibody application above succeeds specifically on the evolutionarily constrained framework rather than on the hypervariable loops. Third, BFC recovers the relative ordering and normalized profile of conformational heterogeneity rather than its absolute magnitude in Ångström units, as reflected in our reliance on rank- and shape-based metrics. Fourth, our validation is entirely computational, resting on structural ensembles, MD trajectories, and docking or refinement benchmarks; direct experimental confirmation and the establishment of functional causality remain to be carried out.

The requirement for an experimental input is not fundamental. Coupling BFC to a front-end that infers B-factors from sequence would in principle extend the same correction to any protein, including those with no deposited structure. We also deliberately withheld three-dimensional coordinates from the model, a controlled design that isolates the contribution of sequence context and demonstrates that a one-dimensional sequence channel alone already carries a decodable record of conformational heterogeneity. A natural next step is to fuse explicit structural information, for example through graph neural networks over the input coordinates, which we expect to raise absolute accuracy and to capture inter-chain rigidification in complexes that a single chain cannot express. Finally, extending BFC to intrinsically disordered proteins, coupling it to wet-lab measurements, and dissecting the mechanistic basis of the conformational heterogeneity it encodes are promising avenues toward turning this retrospective signal into a predictive, causal account of protein motion.

This finding enables dynamic mining of the PDB, allowing researchers to revisit over 190,000 static structures and extract quantitative conformational-heterogeneity information without the prohibitive computational cost of molecular dynamics simulations (Jung et al., 2019; Sahil et al., 2023; Prašnikar et al., 2024; Lappala, 2024; Cui et al., 2025). A compact language model suffices for this translation task, making BFC computationally efficient and deployable for high-throughput screening on standard workstations. While BFC is not a replacement for MD in studying complex transitions, it provides an efficient approximation of intrinsic conformational heterogeneity, serving as a bridge between static structural biology and dynamic biochemistry. We anticipate that BFC will facilitate the large-scale retrospective analysis of PDB entries to investigate mechanisms of protein function and disease.

## Methods

### Dataset Construction and Preprocessing

We constructed a benchmark connecting static PDB structures with conformational-heterogeneity profiles derived from structural ensembles.

We retrieved high-resolution X-ray structures from the Research Collaboratory for Structural Bioinformatics (RCSB) Protein Data Bank (PDB) and matched them with groundtruth conformational-heterogeneity profiles derived from the PDBFlex database (Hrabe et al., 2016). PDBFlex aggregates all depositions of the same protein (identified by a 95% sequence identity threshold), superimposes them, and reports the per-residue average local C*α* RMSD across this structural ensemble. Concretely, for each cluster PDBFlex designates the longest, most complete chain as the representative (*master*) *m*; for a residue *i* the reported value is the average, over the other *N* − 1 cluster members, of the local C*α* RMSD computed within a 10-residue sliding window *W* (*i*) after superposition onto the master:

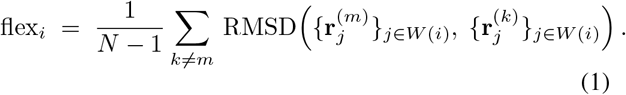

The reference frame is therefore the representative structure rather than the ensemble mean, and the quantity is a member-averaged local deviation rather than an all-pairs (global) average. This per-residue profile is downloaded directly from PDBFlex and used as our ground-truth target without recomputation. It is closely analogous to the Root Mean Square Fluctuation (RMSF) of an ensemble and is hereafter termed *ensemble RMSF* for consistency. This metric quantifies conformational diversity sampled across different crystal forms, ligand states, and refinement conditions, rather than time-resolved dynamics from a single molecular dynamics trajectory. Our training signal therefore reflects the breadth of experimentally observed conformational space, not kinetic fluctuations along a simulation timeline. We consider independent validation against true MD-derived RMSF (see ATLAS validation below) essential for establishing the physical relevance of BFC predictions. To ensure that the model learns from reliable physical signals rather than noise, we applied a strict quality control pipeline: (1) *Experimental Constraints*: Beyond relying on database metadata, we parsed the experimental-method record of each PDB header to confirm X-ray diffraction as the method. Only structures with a resolution ≤ 3.0 Å were retained. (2) *Sequence Processing and Alignment*: A strict residue-level mapping was enforced. Non-standard Selenomethionine (MSE) residues were mapped to Methionine (MET) to handle common crystallographic phasing derivatives, while other non-standard residues were treated as gaps. Chains with significant missing electron density (gaps *>* 10%) or mismatches between C*α* coordinates and the sequence registry were discarded. (3) *Statistical Quality Control*: To filter out corrupted data entries or flat-lined profiles, we calculated the standard deviation of input B-factors and ground-truth RMSF for each chain. Entries exhibiting effectively zero variance (*σ <* 10^*−*6^) were considered artifactual and removed. (4) *Context Sufficiency*: Peptide chains shorter than 50 residues were excluded to ensure sufficient sequence context for the language model. The final curated dataset comprises 280,487 protein chains derived from 132,386 unique PDB entries, partitioned at the PDB-entry level into training (207,465 chains), validation (52,268 chains), and testing (20,754 chains) sets.

Consistent with our goal of decoding intrinsic signals solely from crystallographic data, we minimized the use of hand-crafted features. The model inputs are restricted to two modalities: the raw amino acid sequence (providing physico-chemical context) and the experimental per-residue C*α* B-factors (providing the latent dynamic signal). Because differing refinement protocols and crystal-packing environments give each structure its own B-factor scale, the raw magnitudes are not directly comparable across proteins. We therefore standardized the input B-factors (*B*_*i*_) and target RMSF values (*R*_*i*_) using chain-specific Z-score normalization,

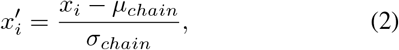

where *µ*_*chain*_ and *σ*_*chain*_ denote the mean and standard deviation for the chain. This normalization compels the model to focus on the relative distribution of fluctuations along the sequence rather than absolute global magnitudes, which are often artifactual.

### Feature Engineering and Input Representation

We restricted the input, excluding explicit structural descriptors derived from coordinate geometry (such as secondary structure assignments, solvent accessibility, or contact maps) and theoretical computations (such as Normal Mode Analysis), so that the model relies strictly on two input modalities. The first is a sequence embedding, in which the raw amino acid sequence of each chain serves as the sole source of physicochemical and evolutionary context. Sequences were tokenized and processed by the pre-trained ESM-2 language model (Lin et al., 2023). We selected the efficient 8-million-parameter variant to keep inference accessible on standard consumer hardware while drawing on the evolutionary features captured by the ESM family, yielding token-level representations h_*i*_ ∈ ℝ ^320^. The second modality is the normalized B-factor, the only experimental observation provided, standardized by the same chain-specific Z-score normalization (Equation (2)) prior to input. The model therefore treats B-factors as noisy proxies for dynamics, using sequence context to denoise and translate these signals into solution-state RMSF.

### B-Factor Corrector (BFC) Architecture

BFC treats dynamics prediction as a token-level regression task. The architecture consists of the ESM-2 encoder followed by a specialized decoding head.

We employed ESM-2 (Lin et al., 2023) as a feature extractor. Unlike approaches that freeze the language model, we performed full-parameter fine-tuning on the ESM-2 backbone. This allows the attention mechanisms to adapt specifically to the task of recognizing crystallographic artifacts and correlating them with sequence motifs.

The embeddings h_*i*_ are concatenated with the normalized B-factor scalar 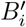. This composite vector is passed through a regression head consisting of a linear projection (dim=256), Layer Normalization, and a ReLU activation function. A Dropout layer (*p* = 0.1) is applied for regularization before the final linear projection to a scalar value representing the predicted RMSF.

### Baseline Models

We benchmarked BFC against three distinct classes of models to isolate the source of performance gains, followed by a sequence-only ablation of the experimental channel.

The first baseline is an analytical formula, a traditional physical conversion in which RMSF is derived directly from B-factors assuming an isotropic harmonic oscillator (Kuzmanic and Zagrovic, 2010),

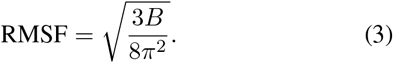

The second baseline is a gradient-boosted decision tree regressor (XGBoost (Chen and Guestrin, 2016)), which tests whether the relationship between B-factors and RMSF is simply a non-linear mapping independent of sequence context; it was trained using *only* the normalized B-factors as input features, with hyperparameters (max depth=5, n estimators=1000) optimized on the validation set. The third baseline is a frozen ESM-2, which assesses the necessity of fine-tuning by training the same prediction head described above on top of fixed embeddings from the pre-trained ESM-2 model, without updating the transformer weights.

Finally, to isolate the specific contribution of the experimental B-factor channel, we trained a sequence-only variant in which the normalized B-factor scalar is removed from the input, so that the regression head operates on the ESM-2 token embeddings alone. This ablation was trained under identical conditions to the full BFC model (same backbone, optimizer, schedule, and 100-epoch budget) to ensure a fair comparison, differing only in the absence of the B-factor channel.

### Training and Optimization

We minimized the Mean Absolute Error (MAE) between the predicted and ground-truth RMSF values. To handle variable sequence lengths within batches, we implemented a masking mechanism in which padding tokens (assigned a placeholder label of -1.0) are strictly excluded from the computation. The loss is calculated as:

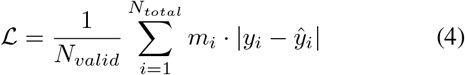

where *m*_*i*_ ∈ {0, 1} is the binary mask indicating valid residues, and *N*_*valid*_ is the total number of non-padding tokens in the batch.

Training was performed using the AdamW optimizer (Loshchilov and Hutter, 2019) with a learning rate of 5×10^*−*5^ and a batch size of 64. We employed a cosine annealing learning rate scheduler with warmup, where the learning rate increases linearly for the first 10% of training steps and then decays following a cosine curve to zero.

All models were implemented in PyTorch 2.6.0 (Paszke et al., 2019) and trained on a single NVIDIA GeForce RTX 5090 (32 GB VRAM, CUDA 13.0). To balance computational efficiency with data integrity, input sequences were truncated to a maximum length of 1,024 residues. Statistical analysis confirms that this threshold covers 99.37% of the training set (Figure S8), ensuring negligible information loss. Training proceeded for a fixed duration of 100 epochs, and the checkpoint achieving the highest Pearson Correlation Coefficient (PCC) on the validation set was selected for testing.

### Evaluation Metrics

We employed four metrics spanning relative fluctuation patterns and absolute physical magnitudes. Because each chain undergoes chain-specific Z-score normalization prior to prediction (removing inter-chain scale differences), all metrics are computed globally over the *M* total residues in the test set. Let y = (*y*_1_, …, *y*_*M*_ ) denote the concatenated predicted values and ŷ = (*ŷ* _1_, …, *ŷ*_*M*_ ) denote the concatenated groundtruth values across all test chains.

The Pearson correlation coefficient (PCC) measures the linear correlation between predicted and experimental values across all residues, assessing whether the model captures the shape of the fluctuation landscape,

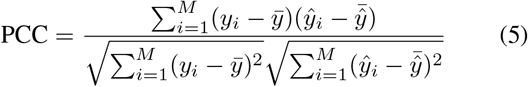

The Spearman correlation coefficient (SCC) evaluates the monotonic rank-order correlation. This metric is robust to outliers and assesses whether the model correctly ranks residues from most rigid to most flexible,

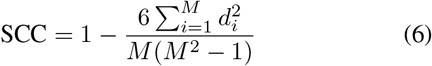

where *d*_*i*_ is the difference between the ranks of *y*_*i*_ and *ŷ*_*i*_.

The root mean square error (RMSE) quantifies the average magnitude of error in physical units (Å ), penalizing large deviations more heavily than small ones,

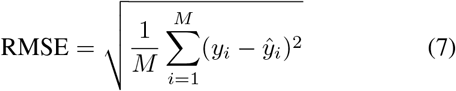

The mean absolute error (MAE) provides a linear score of error magnitude, offering a direct interpretation of the average deviation per residue,

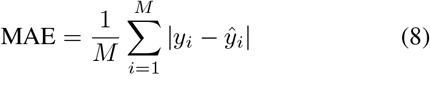

We also assessed physical-space reconstruction through an oracle rescaling strategy. Since solution-state fluctuation amplitudes vary independently of sequence due to experimental conditions (e.g., temperature, crystal quality), direct prediction of absolute RMSF magnitudes from static coordinates is ill-posed. This strategy decouples intrinsic profile accuracy from global amplitude scaling for the physical-space evaluation presented in Figure 1C. For each protein chain, both the BFC output (Z-score) and the baseline physical estimates derived from B-factors were linearly transformed to match the mean (*µ*_*GT*_ ) and standard deviation (*σ*_*GT*_ ) of the groundtruth ensemble RMSF. This transformation ensures that the evaluation metrics (RMSE, PCC, SCC) strictly measure the fidelity of the fluctuation landscape shape (profile reconstruction) rather than systematic scaling errors.

### External Validation on ATLAS

To benchmark the biological fidelity of BFC predictions, we utilized the ATLAS database (Vander Meersche et al., 2024) of standardized molecular dynamics simulations. Groundtruth RMSF profiles were established by computing the ensemble average of the three independent trajectory replicates provided for each entry, ensuring robustness against stochastic sampling errors.

For the benchmark-scale analysis (Figures 4 and 5), we used the full set of 1,938 proteins from ATLAS. Because the reference quantities are computed from MD trajectories (a physical observable distinct from the cross-crystal-form ensemble RMSF used for training, and never seen during training), the MD data provide an orthogonal ground truth of a distinct physical class. Each ATLAS entry provides three 100 ns all-atom explicit-solvent replicas (CHARMM36m force field (Huang et al., 2017), frames every 10 ps). Two of the ground-truth channels were taken directly from the ATLAS release, namely per-residue trajectory RMSF and the entropy-based index *N*_eq_ (the effective number of Protein Blocks sampled per residue, a local backbone structural-alphabet diversity computed by ATLAS with PBxplore (Barnoud et al., 2017)), as was AlphaFold2 pLDDT (Jumper et al., 2021), which ATLAS distributes as a per-residue prediction-confidence score. Because pLDDT quantifies the confidence of the model in its predicted co-ordinates rather than physical flexibility (Vander Meersche et al., 2025a), we treat it throughout only as a heuristic uncertainty proxy (a sequence-derived comparator, not an MD observable) for which low prediction confidence empirically coincides with poorly ordered, mobile regions (Guo et al., 2022) (sign-inverted so that low confidence maps to high predicted flexibility). The remaining quantities were computed from the raw ATLAS trajectories: backbone dihedral entropy *S*_*φψ*_ (Shannon entropy of the joint *φ/ψ* distribution on a 36 × 36 grid, per replica then averaged); a per-residue interaction-energy fluctuation *σ*_*E*_ (standard deviation over frames of the CHARMM36 nonbonded energy, evaluated with an OpenMM CHARMM36 parameterization (Eastman et al., 2024), summed over neighboring residues within 1.2 nm); a conformational-state transition rate (each residue’s backbone (*φ, ψ*) assigned per frame to one of three Ramachandran basins, with boundaries placed in low-density regions of the (*φ, ψ*) map so that a boundary crossing corresponds to traversing a genuine barrier rather than to thermal jitter within a basin: *α* for *φ <* 0^*°*^ and −120^*°*^ *< ψ <* 50^*°*^, *β*/PPII for *φ <* 0^*°*^ otherwise, and *α*_*L*_ for the sparsely populated positive-*φ* region *φ* ≥ 0^*°*^; a ≥ 5-frame (≥ 50 ps) dwell filter was then applied so that a basin change is accepted only when the new basin persists for at least five consecutive frames, shorter excursions being merged back into the preceding stable state, and confirmed transitions were counted *within each replica separately* and then averaged; no transition is counted across a replica boundary); and an intra/interstate fluctuation decomposition. Global conformational states were defined from a C*α*-PCA free-energy landscape: perframe C*α* coordinates were mean-centered, projected onto their first two principal components, binned on a 40 × 40 grid to give an occupancy *P* and a free energy *F* = − ln *P* (in units of *k*_*B*_*T*, shifted to min *F* = 0); each connected region with *F <* 1 *k*_*B*_*T* was taken as a basin, and every frame was assigned to its nearest basin centre. The total per-residue mean-square fluctuation was partitioned via the law of total variance into a within-basin (intra-state) and a between-basin (inter-state) component,

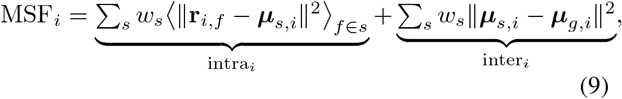

where *w*_*s*_ is the occupancy fraction of state *s, µ*_*s,i*_ the mean position of residue *i* within state *s*, and *µ*_*g,i*_ its global mean position. We note that *N*_eq_ (Protein Blocks) and *S*_*φψ*_ (*φ/ψ*) both quantify local backbone dihedral diversity and are correlated (per-protein median Spearman 0.62); we therefore treat their agreement as robustness across an independent pipeline rather than as statistically independent cross-validation. BFC profiles and all channels were per-chain z-scored and aligned by author residue numbering. Peaks were called at a prominence threshold of ≥ 1*σ*, and two co-localization statistics were computed against each channel across top-*N* % thresholds (*N* = 5–25). (i) *Symmetric-Jaccard region overlap*: taking the top-*N* % residues of the BFC and channel profiles within each chain as high-region sets *A* and *B*, we report the pooled Jaccard index *J* = _*c*_ |*A*_*c*_ ∩ *B*_*c*_|*/* _*c*_ |*A*_*c*_ ∪ *B*_*c*_| over chains *c*; it is symmetric because both sets use the same top-*N* % cut. (ii) *Peak co-localization* (peak precision): the fraction of BFC peak apices falling within the channel’s top-*N* % high region. Significance was assessed against a within-chain permutation null (NPERM = 5000): within each chain the BFC high-region residues (or peak positions) were randomly re-placed, preserving their count and the chain length and channel distribution, with the channel held fixed, and the statistic recomputed; enrichment is the observed value over the null median, and *p* = (#{null ≥ obs} + 1)*/*(NPERM + 1). A shuffled-BFC profile served as an additional negative control; the probability-calibration analysis (Figure 5B) instead binned residues by BFC score and evaluated the monotonic trend against the same null. Termini (five residues per chain end) were excluded from the switching analysis to avoid fraying artifacts, leaving 1,930 proteins. The intra/inter decomposition is only well-posed where conformational states are resolved: in the full multibasin set the two components are collinear (per-protein median correlation 0.75), so we restricted this specific analysis to the 592 proteins with a clean inter-basin separation. For each landscape, the barrier between its two deepest basins was measured by progressively raising a flood-fill threshold until the two basin minima first became connected and taking the height of that connecting level above the deeper minimum; proteins with two or more basins and a barrier ≥ 2.5 *k*_*B*_*T* (at 300 K) were retained. We further tested generalization on 118 apo–holo protein pairs curated from CryptoSite (Cimermancic et al., 2016) and PocketMiner (Meller et al., 2023); for each pair the per-residue conformational change from apo to holo was quantified as 1−(local CAD-score (Olechnovič et al., 2013)), and BFC (predicted from the apo structure and B-factors alone, without ever seeing the bound state) was tested for co-localization with the high-change residues by the same region-overlap and peak-precision statistics against a within-chain permutation null. Exact statistics for all analyses are tabulated in Tables S3 to S6.

### Conformational Ensemble Comparison

To position BFC relative to deep-learning ensemble generators (Figure S4), we generated conformational ensembles for 300 held-out proteins. These comprise 87 proteins in the intersection of the ATLAS set (providing MD-derived RMSF) and the BFC held-out test split (which additionally provides PDBFlex (Hrabe et al., 2016) ensemble RMSF as a ground truth), together with 213 further proteins drawn at random from ATLAS to extend the generator-versus-MD comparison. Analyses that require the PDBFlex reference (the five-way correlation structure and the residue-class contrasts) are therefore restricted to the 87-protein subset, which is statistically representative of the full ATLAS set in both chain length and MD-fluctuation magnitude (two-sample Kolmogorov–Smirnov test, *p >* 0.5 for both). The generator–MD and BFC–MD correlations, which do not need PDBFlex, use all 300 proteins. For each protein we generated a conformational ensemble with two ensemble generators run from the single query sequence: ESMFlow using the PDB-trained distilled checkpoint (ESMFold backbone, run in its non-diffusion sampling mode from a noisy initial state, 80 conformers per protein) (Jing et al., 2024), and BioEmu (AlphaFold2 weights, 80 samples per protein) (Lewis et al., 2025). The generator we used is *ESMFlow-PDB*: the variant of the AlphaFlow/ESMFlow framework fine-tuned on Protein Data Bank structures to model experimental ensembles (the conformational states in which the same protein is deposited across different X-ray crystallography or cryo-EM conditions) as opposed to its molecular-dynamics-trained counterpart (ESMFlow-MD/AlphaFlow-MD) fitted to ATLAS trajectories at 300 K (Jing et al., 2024). We selected this variant precisely because it is the member of the family trained on cross-deposition structural heterogeneity (the observable that BFC targets), so that the comparison is made against the generator best positioned to reproduce it. From each generated ensemble we computed a per-residue fluctuation profile as the C*α* root-mean-square fluctuation about the mean position after rigid-body superposition,

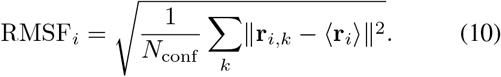

The same RMSF estimator was applied to the ATLAS MD trajectories, placing both coordinate ensembles on a common footing; the PDBFlex reference retains its database definition as the per-residue average local C*α* RMSD across depositions (Hrabe et al., 2016). As an additional physics-based, sequence-blind baseline we computed anisotropic network model (ANM) square fluctuations from each single input structure with ProDy (C*α* network, cutoff 15 Å, lowest 40 non-trivial modes) (Atilgan et al., 2001). All profiles (BFC, generated-ensemble RMSF, ANM, experimental B-factor, ATLAS MD-RMSF, and PDBFlex ensemble RMSF) were aligned by residue numbering, per-chain *z*-scored, and compared by per-protein Spearman correlation (reported as the median across the 87 proteins) so that between-chain amplitude variance is not conflated with within-chain profile agreement. To isolate the residues that discriminate the two axes, we labeled each residue from its within-chain *z*scores by a top-quartile rule: *heterogeneity-specific* if its PDBFlex ensemble-RMSF ranked in the top 25% within its chain while its MD fluctuation did not, *thermal-specific* in the converse case (MD in the top 25%, PDBFlex not), with residues high or low on both axes excluded from this contrast; we tabulated its structural context (relative Solvent Accessible Surface Area, SASA, and DSSP secondary structure), and compared per-method scores on the two classes by a per-protein paired Wilcoxon signed-rank test. Finally, on strictly held-out test proteins we binned residues into tertiles by input B-factor and by BFC prediction (both within-protein *z*) and reported the mean ground-truth PDBFlex heterogeneity per cell, with 95% confidence intervals from a protein-block bootstrap (*B* = 5,000).

### Functional Annotation

For the functional case studies presented (e.g., PDB 1D4T and 1CUO), critical functional regions such as ligand-binding loops and active-site adjacencies were identified and mapped according to the UniProt Knowledgebase (UniProt Consortium, 2018) annotations. To quantitatively verify the correspondence between predicted dynamics and biological function, we implemented a statistical scoring workflow. The BFC-predicted normalized fluctuations of these UniProt-annotated regions were compared against the remaining structural scaffold. Welch’s t-test for unequal variances was used to assess significance, with a threshold of *p <* 0.001 confirming that the model selectively assigns significantly higher flexibility to function-critical motifs compared to the stable protein core.

### Sparse Autoencoder Interpretability Analysis

To examine how BFC internally represents conformational heterogeneity, we performed a post-hoc analysis of its layer-6 activations with a sparse autoencoder (SAE), which decomposes the dense representation into an interpretable feature dictionary. This analysis asks whether the heterogeneity signal that BFC is trained to recover is organized as a sparse, separable set of features rather than diffusely entangled across the representation.

For each residue we extracted the final-layer (layer-6) representation of the fine-tuned BFC backbone (ESM-2, dimension *D* = 320) across the full training corpus (56,791,414 residues). We used a Top-*K* SAE (Gao et al., 2025): an encoder followed by a hard top-*K* selection (only the *K* largest activations are retained and passed through a ReLU, the rest set to zero) and a linear decoder,

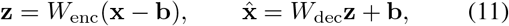

where b is a pre-bias initialized to the data mean. Sparsity is enforced architecturally by the top-*K* constraint alone; no *L*_1_ penalty is applied. Each model was trained to minimize the reconstruction mean squared error in per-dimension standardized space using Adam (learning rate 10^*−*3^, batch size 16,384), halting once the reconstruction quality plateaued and retaining the best-reconstruction checkpoint.

The two hyperparameters governing an SAE are the dictionary size *F* (number of features) and the sparsity level *K* (active features per residue), which trade reconstruction fidelity against feature disentanglement. We treated reconstruction quality and dictionary health as admissibility constraints; a configuration must reconstruct the representation faithfully (high variance explained) without collapsing into unused units (low dead-feature fraction). The number of monosemantic features was the selection objective. Here a monosemantic feature is one whose activation, computed over all 56.8M residues, attains an absolute Pearson correlation with the ensemble RMSF above a fixed threshold (|PCC| *>* 0.2), so that the count reflects how cleanly the dictionary factors this axis into dedicated units.

Rather than exhaustively filling a grid over (*F, K*), we located the optimum with two orthogonal one-dimensional sweeps through the candidate configuration. Holding *F* = 2560 and increasing *K* (32 → 64 → 128) revealed an interior maximum in the monosemantic-feature count: it rose from 79 to 131 and then fell to 89, because too few active units underfit the representation while too many force individual features to absorb several unrelated meanings. Holding *K* fixed and doubling the dictionary width (*F* =2560 → 5120) did not increase the count (from 131 to 99 at *K*=64; from 89 to 82 at *K*=128), indicating that added capacity is spent on entangled, over-complete units rather than on new interpretable features. The remaining cells of the grid lie strictly in these already-worse directions and were therefore not evaluated. The selected configuration (*F* = 2560, *K* = 64) keeps reconstruction high (variance explained 0.87) with negligible dead features (0.2%) while yielding the largest number of monosemantic features. Although the higher-capacity configurations (*K*=128) reach a higher variance explained (≈ 0.95), they do so through over-complete dictionaries that resolve fewer interpretable features. The selection is stable across the relevant range. The chosen configuration yields the most monosemantic features at every threshold from 0.15 to 0.30, and only at the most permissive threshold (0.10) do the over-complete configurations lead, where weak noise-inflated correlations dominate the count (Figure S5).

To characterize individual features (Figure 8), we defined the top activating sites of a feature as the residues whose activation exceeds the 90th percentile of that feature’s non-zero activations. At these sites we tabulated amino-acid composition, secondary structure (DSSP), relative solvent accessibility, and the percentile ranks of the input B-factor and ensemble RMSF, and we recomputed the feature–heterogeneity correlation within each secondary-structure class to confirm that the association is not a trivial consequence of secondary structure.

### Molecular Docking Benchmark

To establish a rigorous paired benchmark connecting sequence-derived dynamics with binding conformational changes, we curated the PDBbind Refined Set (v2020) (Wang et al., 2005; Liu et al., 2017), comprising 5,316 entries. From this initial pool, we applied a strict multi-stage filtration pipeline to ensure chemical and structural consistency. Complexes exhibiting missing receptor backbones, inconsistent ligand chemistry, or failures in geometric center calculation during Meeko preparation (Santos-Martins et al., 2025) were excluded. To maintain a strictly paired statistical design, we retained only those targets that successfully completed all three parallel docking protocols (Rigid, Experimental-guided, and BFC-guided). This intersection reduced the initial dataset to a benchmark of 1,204 protein-ligand complexes, so that performance metrics reflect algorithmic differences rather than dataset heterogeneity.

We utilized the fine-tuned BFC model to infer residue-level flexibility for each target chain. The amino acid sequence was tokenized and processed by the ESM-2 backbone, and the resulting latent representations were combined with the per-residue experimental C*α* B-factors and decoded into denoised normalized flexibility scores (Z-scores). This inference therefore operated on sequence context together with experimental B-factors, without requiring three-dimensional coordinate topology or contact maps; the role of sequence context is to decouple intrinsic flexibility from crystal-packing artifacts embedded in the experimental B-factor signal.

To evaluate the physical fidelity of the predicted dynamics, we implemented a controlled budget-comparison strategy. The binding pocket was defined as all residues within a 6.0 Å radius of the crystal ligand. We compared two distinct flexibility selection protocols against a rigid baseline. In the experimental protocol, the three residues within the pocket exhibiting the highest crystallographic C*α* B-factors were designated as flexible. In the BFC protocol, the top three residues with the highest predicted flexibility scores were selected. By fixing the degree-of-freedom budget at *N* = 3 residues, we ensured that any divergence in docking performance was attributable solely to the quality of the flexibility identification source rather than the complexity of the conformational search space.

Molecular docking simulations were executed using UniDock (Yu et al., 2023), a GPU-accelerated implementation of the AutoDock Vina (Trott and Olson, 2010) scoring function. To ensure robust sampling of the expanded conformational landscape induced by side-chain flexibility, we employed a high exhaustiveness setting of 128 with a maximum of 20 generation steps. The search space was dynamically defined for each target, centered on the native ligand geometry with a padding of 10.0 Å in all dimensions. Simulations were executed in isolated threads to enforce deterministic reproducibility. Docking accuracy was quantified by the Root Mean Square Deviation (RMSD) of the top-ranked pose relative to the crystal reference, with a success threshold defined at ≤ 2.0 Å . To compare the two flexibility sources across the benchmark, we plotted cumulative success curves (the fraction of targets with pose RMSD below a sweep of thresholds) for the rigid, experimental-B-factor, and BFC protocols. Statistical equivalence between the BFC-guided and experimental-B-factor RMSD distributions was assessed by two one-sided tests (TOST) (Schuirmann, 1987) at a 0.5 Å equivalence margin, reporting the equivalence *P* value and the mean paired difference.

### Antibody Interface Refinement

A diverse dataset of 30 antibody-antigen complexes was curated from SAbDab (Dunbar et al., 2014). Initial singlechain variable fragment (scFv) structures were predicted using ESMFold with a flexible linker (Lin et al., 2023). These predictions were superposed onto the native antibody coordinates relative to the antigen using heavy-chain framework C*α* atoms, which fixes the binding pose from the known complex and deliberately preserves the interface steric clashes introduced by rigid-body placement of an unliganded model. This constitutes a controlled interface-refinement setup; the objective is to relieve these clashes by local induced fit, not to perform blind docking or to search for the binding orientation. Because ESMFold models carry no experimental temperature factors, the per-residue pLDDT confidence was used in place of the B-factor scalar channel; as lower pLDDT corresponds to locally less certain, typically more flexible regions, it was inverted as 100−pLDDT so that low confidence maps to high flexibility, and Z-score normalized per chain before input.

Refinement was conducted using OpenMM (v8.4) (Eastman et al., 2024) with the Amber ff14SB force field (Maier et al., 2015) and the GB-Neck2 implicit solvent model (Nguyen et al., 2013). Pre-processing included pH 7.0 protonation, automatic disulfide bond reconstruction (CYS to CYX), and terminal oxygen geometry correction. We implemented a differential restraint scheme based on the BFC flexibility map (threshold *τ* = 0.5). Harmonic positional restraints were applied to the C*α* atom of each residue, tethering it to its initial coordinate: strict restraints held the antigen (*k* = 500 kJ/mol/nm^2^) and the rigid antibody frameworks (*k* = 100 kJ/mol/nm^2^) to maintain topology, while flexible loops received minimal restraints (*k* = 1.0 kJ/mol/nm^2^) to enable induced fit. The protocol employed an adaptive Langevin dynamics strategy: following multi-stage energy minimization, a 100 ps production run was performed at 300 K. An automated fallback mechanism activated a stable mode (reduced time step 0.5 fs, high friction 5.0 ps^*−*1^) for strained starting structures that would otherwise be numerically unstable. To attribute refinement quality to the BFC prior rather than the MD engine, two further schemes were run on the identical protocol: an expert-annotation control, in which the flexible set was the CDR loops defined by ANARCI Chothia numbering (Dunbar and Deane, 2016) (framework restrained, CDR released), and an unrestrained control (no antibody restraints; antigen fixed). Framework preservation was quantified as the framework-C*α* RMSD from the initial structure after superposition on the framework (drift from input), which removes the constant ESMFold prediction error; the interface clash count was defined as the number of antibody heavy atoms within 2.0 Å of any antigen heavy atom. Paired schemes were compared over the 30 complexes by two-sided Wilcoxon signed-rank tests. Representative before/after conformations (PDB 9EHT) were rendered in PyMOL.

## Data and code availability

Source code for BFC and the pre-trained model weights are freely available at https://github.com/wyqmath/BFactorCorrector. All datasets analyzed in this study are publicly available from their original repositories without restriction.

## References

Atilgan, A.R., Durell, S.R., Jernigan, R.L., Demirel, M.C., Keskin, O., and Bahar, I. (2001). Anisotropy of fluctuation dynamics of proteins with an elastic network model. Biophys J 80, 505–515.

Aurora, R., Srinivasan, R., and Rose, G.D. (1994). Rules for α-helix termination by glycine. Science 264, 1126–1130.

Bahar, I., Atilgan, A.R., and Erman, B. (1997). Direct evaluation of thermal fluctuations in proteins using a single-parameter harmonic potential. Fold Des 2, 173–181.

Barnoud, J., Santuz, H., Craveur, P., Joseph, A.P., Jallu, V., de Brevern, A.G., and Poulain, P. (2017). Pbxplore: a tool to analyze local protein structure and deformability with protein blocks. PeerJ 5, e4013.

Berman, H.M., Westbrook, J., Feng, Z., Gilliland, G., Bhat, T.N., Weissig, H., Shindyalov, I.N., and Bourne, P.E. (2000). The protein data bank. Nucleic Acids Res 28, 235– 242.

Bhakat, S., and Strauch, E.M. (2026). Accelerated sampling of protein dynamics using BioEmu-augmented molecular simulation. J Chem Inf Model 66, 7168–7178. doi: 10.1021/acs.jcim.6c01000.

Blackler, R.J., Müller-Loennies, S., Pokorny-Lehrer, B., Legg, M.S., Brade, L., Brade, H., Kosma, P., and Evans, S.V. (2022). Antigen binding by conformational selection in near-germline antibodies. J Biol Chem 298, 101901.

Booth, C., Gilmour, K.C., Veys, P., Gennery, A.R., Slatter, M.A., Chapel, H., and Gaspar, H.B. (2011). X-linked lymphoproliferative disease due to SAP/SH2D1A deficiency: a multicenter study on the manifestations, management and outcome of the disease. Blood 117, 53–62.

Carugo, O. (2022). B-factor accuracy in protein crystal structures. Acta Crystallogr D Struct Biol 78, 69–74.

Chakravarty, D., Schafer, J.W., Chen, E.A., Thole, J.F., Ronish, L.A., Lee, M., and Porter, L.L. (2024). AlphaFold predictions of fold-switched conformations are driven by structure memorization. Nat Commun 15, 7296.

Chen, T., and Guestrin, C. XGBoost: A scalable tree boosting system. In Proceedings of the 22nd ACM SIGKDD International Conference on Knowledge Discovery and Data Mining, KDD ‘16, pages 785–794, New York, NY, USA, (2016). Association for Computing Machinery. ISBN 9781450342322.

Chou, P.Y., and Fasman, G.D. (1974). Prediction of protein conformation. Biochemistry 13, 222–245.

Cilia, E., Pancsa, R., Tompa, P., Lenaerts, T., and Vranken, W.F. (2014). The DynaMine webserver: predicting protein dynamics from sequence. Nucleic Acids Res 42, W264– W270.

Cimermancic, P., Weinkam, P., Rettenmaier, T.J., Bichmann, L., Keedy, D.A., Woldeyes, R.A., Schneidman-Duhovny, D., Demerdash, O.N., Mitchell, J.C., Wells, J.A., et al. (2016). CryptoSite: expanding the druggable proteome by characterization and prediction of cryptic binding sites. J Mol Biol 428, 709–719.

Csermely, P., Palotai, R., and Nussinov, R. (2010). Induced fit, conformational selection and independent dynamic segments: an extended view of binding events. Trends Biochem Sci 35, 539–546.

Cui, T., Zhou, Y., and Wang, T. (2025). Recent advances in artificial intelligence–driven biomolecular dynamics simulations based on machine learning force fields. Curr Opin Struct Biol 95, 103191.

Dauparas, J., Anishchenko, I., Bennett, N., Bai, H., Ragotte, R.J., Milles, L.F., Wicky, B.I., Courbet, A., de Haas, R.J., Bethel, N., et al. (2022). Robust deep learning–based protein sequence design using ProteinMPNN. Science 378, 49–56.

Dauparas, J., Lee, G.R., Pecoraro, R., An, L., Anishchenko, I., Glasscock, C., and Baker, D. (2025). Atomic context-conditioned protein sequence design using LigandMPNN. Nat Methods pages 1–7.

DePristo, M.A., De Bakker, P.I., and Blundell, T.L. (2004). Heterogeneity and inaccuracy in protein structures solved by X-ray crystallography. Structure 12, 831–838.

Dunbar, J., and Deane, C.M. (2016). ANARCI: antigen receptor numbering and receptor classification. Bioinformatics 32, 298–300.

Dunbar, J., Krawczyk, K., Leem, J., Baker, T., Fuchs, A., Georges, G., Shi, J., and Deane, C.M. (2014). SAbDab: the structural antibody database. Nucleic Acids Res 42, D1140–D1146.

Eastman, P., Galvelis, R., Peláez, R.P., Abreu, C.R., Farr, S.E., Gallicchio, E., Gorenko, A., Henry, M.M., Hu, F., Huang, J., et al. (2024). OpenMM 8: molecular dynamics simulation with machine learning potentials. J Phys Chem B 128, 109–116.

Eraslan, G., Simon, L.M., Mircea, M., Mueller, N.S., and Theis, F.J. (2019). Single-cell RNA-seq denoising using a deep count autoencoder. Nat Commun 10, 390.

Eyal, E., Gerzon, S., Potapov, V., Edelman, M., and Sobolev, V. (2005). The limit of accuracy of protein modeling: influence of crystal packing on protein structure. J Mol Biol 351, 431–442.

Fenwick, R.B., van den Bedem, H., Fraser, J.S., and Wright, P.E. (2014). Integrated description of protein dynamics from room-temperature X-ray crystallography and NMR. Proc Natl Acad Sci 111, E445–E454.

Fernández-Quintero, M.L., Kraml, J., Georges, G., and Liedl, K.R. (2019). CDR-H3 loop ensemble in solution–conformational selection upon antibody binding. MAbs 11, 1077–1088.

Fernández-Quintero, M.L., Georges, G., Varga, J.M., and Liedl, K.R. (2021). Ensembles in solution as a new paradigm for antibody structure prediction and design. MAbs 13, 1923122.

Frauenfelder, H., Sligar, S.G., and Wolynes, P.G. (1991). The energy landscapes and motions of proteins. Science 254, 1598–1603.

Gao, L., Dupre la Tour, T., Tillman, H., Goh, G., Troll, R., Radford, A., Sutskever, I., Leike, J., and Wu, J. Scaling and evaluating sparse autoencoders. In International Conference on Learning Representations, volume 2025, pages 26721–26754, (2025).

Guo, H.B., Perminov, A., Bekele, S., Kedziora, G., Farajollahi, S., Varaljay, V., Hinkle, K., Molinero, V., Meister, K., Hung, C., et al. (2022). AlphaFold2 models indicate that protein sequence determines both structure and dynamics. Sci Rep 12, 10696.

Hauser, T., Bhat, J.Y., Miličić, G., Wendler, P., Hartl, F.U., Bracher, A., and Hayer-Hartl, M. (2015). Structure and mechanism of the Rubisco-assembly chaperone Raf1. Nat Struct Mol Biol 22, 720–728.

Hayes, T., Rao, R., Akin, H., Sofroniew, N.J., Oktay, D., Lin, Z., Verkuil, R., Tran, V.Q., Deaton, J., Wiggert, M., et al. (2025). Simulating 500 million years of evolution with a language model. Science 387, 850–858.

Högel, P., Götz, A., Kuhne, F., Ebert, M., Stelzer, W., Rand, K.D., Scharnagl, C., and Langosch, D. (2018). Glycine perturbs local and global conformational flexibility of a transmembrane helix. Biochemistry 57, 1326–1337.

Hou, C., Zhao, H., and Shen, Y. (2026). Protein language models trained on biophysical dynamics inform mutation effects. Proc Natl Acad Sci 123, e2530466123.

Hrabe, T., Li, Z., Sedova, M., Rotkiewicz, P., Jaroszewski, L., and Godzik, A. (2016). PDBFlex: exploring flexibility in protein structures. Nucleic Acids Res 44, D423–D428.

Hu, G., Katuwawala, A., Wang, K., Wu, Z., Ghadermarzi, S., Gao, J., and Kurgan, L. (2021). flDPnn: Accurate intrinsic disorder prediction with putative propensities of disorder functions. Nat Commun 12, 4438.

Huang, J., Rauscher, S., Nawrocki, G., Ran, T., Feig, M., De Groot, B.L., Grubmüller, H., and MacKerell Jr, A.D. (2017). CHARMM36m: an improved force field for folded and intrinsically disordered proteins. Nat Methods 14, 71– 73.

Huben, R., Cunningham, H., Smith, L., Ewart, A., and Sharkey, L. Sparse autoencoders find highly interpretable features in language models. In International Conference on Learning Representations, volume 2024, pages 7827– 7845, (2024).

Ilzhöfer, D., Heinzinger, M., and Rost, B. (2022). SETH predicts nuances of residue disorder from protein embeddings. Front Bioinform 2, 1019597.

Inoue, T., Suzuki, S., Nishio, N., Yamaguchi, K., Kataoka, K., Tobari, J., Yong, X., Hamanaka, S., Matsumura, H., and Kai, Y. (2003). The significance of the flexible loop in the azurin (Az-iso2) from the obligate methylotroph Methylomonas sp. strain j. J Mol Biol 333, 117–124.

Jing, B., Berger, B., and Jaakkola, T. AlphaFold meets flow matching for generating protein ensembles. In Forty-first International Conference on Machine Learning, (2024).

Jumper, J., Evans, R., Pritzel, A., Green, T., Figurnov, M., Ronneberger, O., Tunyasuvunakool, K., Bates, R., Žídek, A., Potapenko, A., et al. (2021). Highly accurate protein structure prediction with AlphaFold. Nature 596, 583–589.

Jung, J., Nishima, W., Daniels, M., Bascom, G., Kobayashi, C., Adedoyin, A., and Sanbonmatsu, K.Y. (2019). Scaling molecular dynamics beyond 100,000 processor cores for large-scale biophysical simulations. J Comput Chem 40, 1919–1930.

Karplus, M., and McCammon, J.A. (2002). Molecular dynamics simulations of biomolecules. Nat Struct Biol 9, 646–652.

Karplus, P.A. (2015). A primer on peroxiredoxin biochemistry. Free Radic Biol Med 80, 183–190.

Klyshko, E., Kim, J.S.H., McGough, L., Valeeva, V., Lee, E., Ranganathan, R., and Rauscher, S. (2024). Functional protein dynamics in a crystal. Nat Commun 15, 3244.

Krieger, F., Möglich, A., and Kiefhaber, T. (2005). Effect of proline and glycine residues on dynamics and barriers of loop formation in polypeptide chains. J Am Chem Soc 127, 3346–3352.

Kuzmanic, A., and Zagrovic, B. (2010). Determination of ensemble-average pairwise root mean-square deviation from experimental B-factors. Biophys J 98, 861–871.

Lappala, A. (2024). The next revolution in computational simulations: Harnessing AI and quantum computing in molecular dynamics. Curr Opin Struct Biol 89, 102919.

Lasorsa, A., and van Der Wel, P.C. (2025). Solid-state NMR protocols for unveiling dynamics and (drug) interactions of membrane-bound proteins. Protein Sci 34, e70102.

Lewis, S., Hempel, T., Jiménez-Luna, J., Gastegger, M., Xie, Y., Foong, A.Y.K., Satorras, V.G., Abdin, O., Veeling, B.S., Zaporozhets, I., et al. (2025). Scalable emulation of protein equilibrium ensembles with generative deep learning. Science 389, eadv9817.

Li, D.W., and Brüschweiler, R. (2009). All-atom contact model for understanding protein dynamics from crystallographic B-factors. Biophys J 96, 3074–3081.

Lin, Z., Akin, H., Rao, R., Hie, B., Zhu, Z., Lu, W., Smetanin, N., Verkuil, R., Kabeli, O., Shmueli, Y., et al. (2023). Evolutionary-scale prediction of atomic-level protein structure with a language model. Science 379, 1123– 1130.

Liu, C.N., Denzler, L.M., Hood, O.E., and Martin, A.C. (2024). Do antibody CDR loops change conformation upon binding? MAbs 16, 2322533.

Liu, Z., Su, M., Han, L., Liu, J., Yang, Q., Li, Y., and Wang, R. (2017). Forging the basis for developing protein–ligand interaction scoring functions. Acc Chem Res 50, 302–309.

Loshchilov, I., and Hutter, F. Decoupled weight decay regularization. In International Conference on Learning Representations, (2019).

Maddipatla, A., Sellam Bojan, N., Bojan, M., Masalitin, V., Vedula, S., Schanda, P., Marx, A., and Bronstein, A.M. (2026). Experiment-guided AlphaFold3 resolves measurement-consistent protein ensembles. Nat Biotechnol pages 1–9.

Maier, J.A., Martinez, C., Kasavajhala, K., Wickstrom, L., Hauser, K.E., and Simmerling, C. (2015). ff14sb: improving the accuracy of protein side chain and backbone parameters from ff99sb. J Chem Theory Comput 11, 3696–3713.

Meller, A., Ward, M., Borowsky, J., Kshirsagar, M., Lotthammer, J.M., Oviedo, F., Ferres, J.L., and Bowman, G.R. (2023). Predicting locations of cryptic pockets from single protein structures using the PocketMiner graph neural network. Nat Commun 14, 1177.

Mukherjee, S., and Hummer, G. (2026). Bayesian learning of distance metrics beyond RMSD for biomolecule alignment, clustering, and domain identification. arXiv preprint 2607.07352

Nguyen, H., Roe, D.R., and Simmerling, C. (2013). Improved generalized born solvent model parameters for protein simulations. J Chem Theory Comput 9, 2020–2034.

Nussinov, R., Liu, Y., Zhang, W., and Jang, H. (2023). Protein conformational ensembles in function: roles and mechanisms. RSC Chem Biol 4, 850–864.

Olechnovič, K., Kulberkytė, E., and Venclovas, č. (2013). CAD-score: a new contact area difference-based function for evaluation of protein structural models. Proteins 81, 149–162.

Pace, C.N., and Scholtz, J.M. (1998). A helix propensity scale based on experimental studies of peptides and proteins. Biophys J 75, 422–427.

Pandey, A., Liu, E., Graham, J., Chen, W., and Keten, S. (2023). B-factor prediction in proteins using a sequencebased deep learning model. Patterns 4, 100805. doi: 10.1016/j.patter.2023.100805.

Paszke, A., Gross, S., Massa, F., Lerer, A., Bradbury, J., Chanan, G., Killeen, T., Lin, Z., Gimelshein, N., Antiga, L., et al. PyTorch: An imperative style, high-performance deep learning library. In Advances in neural information processing systems, volume 32, (2019).

Poy, F., Yaffe, M.B., Sayos, J., Saxena, K., Morra, M., Sumegi, J., and Eck, M.J. (1999). Crystal structures of the XLP protein SAP reveal a class of SH2 domains with extended, phosphotyrosine-independent sequence recognition. Mol Cell 4, 555–561.

Praşnikar, E., Ljubiė, M., Perdih, A., and Borişek, J. (2024). Machine learning heralding a new development phase in molecular dynamics simulations. Artif Intell Rev 57, 102.

Sahil, M., Sarkar, S., and Mondal, J. (2023). Long-time-step molecular dynamics can retard simulation of protein-ligand recognition process. Biophys J 122, 802–816.

Sanchez-Garcia, R., Gomez-Blanco, J., Cuervo, A., Carazo, J.M., Sorzano, C.O.S., and Vargas, J. (2021). DeepEMhancer: a deep learning solution for cryo-EM volume post-processing. Commun Biol 4, 874.

Santos-Martins, D., He, Y., Eberhardt, J., Sharma, P., Bruciaferri, N., Holcomb, M., Llanos, M.A., Hansel-Harris, A., Barkdull, A.P., Tillack, A.F., et al. (2025). Meeko: Molecule parametrization and software interoperability for docking and beyond. J Chem Inf Model 65, 13045–13050.

Schuirmann, D.J. (1987). A comparison of the two one-sided tests procedure and the power approach for assessing the equivalence of average bioavailability. J Pharmacokinet Biopharm 15, 657–680.

Scott, K.A., Alonso, D.O., Sato, S., Fersht, A.R., and Daggett, V. (2007). Conformational entropy of alanine versus glycine in protein denatured states. Proc Natl Acad Sci 104, 2661–2666.

Song, X., Bao, L., Feng, C., Huang, Q., Zhang, F., Gao, X., and Han, R. (2024). Accurate prediction of protein structural flexibility by deep learning integrating intricate atomic structures and Cryo-EM density information. Nat Commun 15, 5538.

Spoendlin, F.C., Fernández-Quintero, M.L., Raghavan, S.S., Turner, H.L., Gharpure, A., Loeffler, J.R., Wong, W.K., Bujotzek, A., Georges, G., Ward, A.B., et al. (2025). Predicting the conformational flexibility of antibody and T cell receptor complementarity-determining regions. Nat Mach Intell 7, 1755–1767.

Trott, O., and Olson, A.J. (2010). AutoDock Vina: improving the speed and accuracy of docking with a new scoring function, efficient optimization, and multithreading. J Comput Chem 31, 455–461.

UniProt Consortium, T. (2018). UniProt: the universal protein knowledgebase. Nucleic Acids Res 46, 2699–2699.

Uysal, S., Vásquez, V., Tereshko, V., Esaki, K., Fellouse, F.A., Sidhu, S.S., and Kossiakoff, A. (2009). Crystal structure of full-length KcsA in its closed conformation. Proc Natl Acad Sci 106, 6644–6649.

Van Kempen, M., Kim, S.S., Tumescheit, C., Mirdita, M., Lee, J., Gilchrist, C.L., Söding, J., and Steinegger, M. (2024). Fast and accurate protein structure search with Foldseek. Nat Biotechnol 42, 243–246.

Vander Meersche, Y., Cretin, G., Gheeraert, A., Gelly, J.C., and Galochkina, T. (2024). ATLAS: protein flexibility description from atomistic molecular dynamics simulations. Nucleic Acids Res 52, D384–D392.

Vander Meersche, Y., Diharce, J., Gelly, J.C., and Galochkina, T. (2025a). Flexibility or uncertainty? a critical assessment of AlphaFold 2 pLDDT. Structure 33, 2157–2163.

Vander Meersche, Y., Duval, G., Cretin, G., Gheeraert, A., Gelly, J.c., and Galochkina, T. (2025b). PEGASUS: Prediction of MD-derived protein flexibility from sequence. Protein Sci 34, e70221.

Wang, D., Tan, Z., Gao, J., Zhang, S., Shen, J., and Lu, Y. (2025a). Ai4protein: transforming the future of protein design. Science China Life Sciences 68, 2880–2890.

Wang, R., Fang, X., Lu, Y., Yang, C.Y., and Wang, S. (2005). The PDBbind database: methodologies and updates. J Med Chem 48, 4111–4119.

Wang, Y., Cai, M., Dong, Y., Ma, Y., and Wei, K. (2025b). From signal to symphony: Exploring 2D sequence representations for protein function prediction. J Chem Inf Model 65, 12723–12736.

Watson, J.L., Juergens, D., Bennett, N.R., Trippe, B.L., Yim, J., Eisenach, H.E., Ahern, W., Borst, A.J., Ragotte, R.J., Milles, L.F., et al. (2023). De novo design of protein structure and function with RFdiffusion. Nature 620, 1089– 1100.

Xu, H., Li, L., Pan, S., Cheng, P., Wang, Y., Rong, Z., Liu, F., Huang, X., Wang, S., and Shu, W. (2026). Protein foundation models: a comprehensive survey. Science China Life Sciences 69, 1503–1531.

Yang, Y., Lv, Y., Luo, Z., Wang, Q., Xu, G., and Ma, J. (2025). OPUS-BFactor: Predicting protein B-factor with sequence and structure information. Molecules 30, 2570.

Yu, Y., Cai, C., Wang, J., Bo, Z., Zhu, Z., and Zheng, H. (2023). Uni-Dock: GPU-accelerated docking enables ultralarge virtual screening. J Chem Theory Comput 19, 3336–3345.

