## Supplementary Information for "Sequence context decodes multi-state conformational heterogeneity from crystallographic B-factors"

Table S1: **Ablation of the experimental B-factor channel.** We compare the full B-Factor Corrector (BFC), which integrates the protein sequence with the experimental B-factor channel, against a Sequence-Only variant in which this channel is removed from the architecture so that the model operates on sequence alone. Both models are trained identically (see Methods) and evaluated on the full test set ($N = 20{,}754$ chains); the better value in each comparison is shown in bold. We report global metrics (all residues concatenated), the per-protein mean (averaging per-chain correlations), and a per-chain high-flexibility subset comprising the $25\%$ most flexible residues within each chain. The experimental channel improves every metric, and the per-chain Pearson correlation gain is highly significant (one-sided Wilcoxon signed-rank test, $p = 6.3 \times 10^{-13}$). Table S2 resolves this contribution as a function of residue flexibility.

| Metric | BFC | Seq-Only | Δ |
|---|---|---|---|
| Global PCC | **0.832** | 0.823 | +0.009 |
| Global SCC | **0.813** | 0.809 | +0.004 |
| Per-protein mean PCC | **0.780** | 0.770 | +0.010 |
| High-flexibility PCC (top 25%, per-chain) | **0.746** | 0.734 | +0.012 |

Table S2: **Flexibility dependence of the B-factor channel contribution.** To localize where the experimental channel contributes, we pool all test-set residues and stratify them by their ground-truth ensemble RMSF, with *top* $X\%$ denoting the $X\%$ most flexible residues across the dataset. This dataset-wide difficult-subset analysis is orthogonal to the per-chain subset of Table S1: the strata are defined solely by the rank of the ground-truth target, independently of either model. For each stratum we report the full BFC and Sequence-Only models (see Methods) and the relative improvement (Δ%) under Pearson and Spearman correlation, mean absolute error, and root-mean-square error; the better value in each comparison is shown in bold. The benefit of the experimental channel concentrates toward the most flexible residues. Because these narrow strata contain residues of uniformly high flexibility, the restricted variance of the target depresses the absolute correlation coefficients (a range-restriction effect), so we read the trend primarily from the per-residue error metrics, which are immune to this effect. Relative to the sequence-only model, the error reduction more than doubles between all residues and the top $0.5\%$ (MAE, $+1.9\%$ to $+5.2\%$; RMSE, $+2.5\%$ to $+3.2\%$), with a slight attenuation in the most extreme stratum, where a few heavy-tailed residues dominate the absolute error. The correlation gains rise monotonically over the same range, reaching $+14.4\%$ (PCC) and $+10.0\%$ (SCC) at the top $0.5\%$. The stratification target is the ensemble RMSF from PDBFlex, which reflects conformational diversity across crystal forms rather than single-trajectory MD dynamics; the channel is thus most informative in regions of high conformational heterogeneity, where the evolutionary sequence prior alone is least sufficient. Values are reported in normalized space.

| | | PCC | | | SCC | | | MAE | | | RMSE | | |
|---|---|---|---|---|---|---|---|---|---|---|---|---|---|
| Stratum | $N$ | BFC | Seq-Only | Δ% | BFC | Seq-Only | Δ% | BFC | Seq-Only | Δ% | BFC | Seq-Only | Δ% |
| All | 5,701,521 | **0.832** | 0.823 | +1.1 | **0.813** | 0.809 | +0.5 | **0.165** | 0.168 | +1.9 | **0.559** | 0.573 | +2.5 |
| Top 50% | 2,850,762 | **0.819** | 0.809 | +1.3 | **0.654** | 0.647 | +1.1 | **0.252** | 0.258 | +2.1 | **0.748** | 0.767 | +2.5 |
| Top 25% | 1,425,388 | **0.781** | 0.769 | +1.6 | **0.621** | 0.608 | +2.1 | **0.441** | 0.451 | +2.2 | **1.035** | 1.062 | +2.5 |
| Top 10% | 570,153 | **0.649** | 0.631 | +3.0 | **0.565** | 0.547 | +3.1 | **0.804** | 0.828 | +3.0 | **1.571** | 1.613 | +2.6 |
| Top 5% | 285,077 | **0.488** | 0.465 | +5.0 | **0.515** | 0.494 | +4.3 | **1.126** | 1.169 | +3.7 | **2.075** | 2.133 | +2.7 |
| Top 2% | 114,031 | **0.323** | 0.300 | +7.8 | **0.464** | 0.439 | +5.8 | **1.551** | 1.629 | +4.8 | **2.757** | 2.845 | +3.1 |
| Top 1% | 57,016 | **0.252** | 0.233 | +8.0 | **0.436** | 0.412 | +5.8 | **1.866** | 1.979 | +5.7 | **3.253** | 3.371 | +3.5 |
| Top 0.5% | 28,508 | **0.211** | 0.185 | +14.4 | **0.430** | 0.391 | +10.0 | **2.190** | 2.311 | +5.2 | **3.743** | 3.869 | +3.2 |

Table S3: **Co-localization of BFC peaks with MD-derived and sequence-derived disorder channels on the ATLAS benchmark (Figure 4C,D,F,G).** For each channel we report the enrichment of the overlap between BFC high regions and channel high regions, over a pooled within-chain permutation null (NPERM $= 5000$), across five top-$N\%$ thresholds. Two co-localization measures are shown: region overlap (symmetric Jaccard; Figure 4C) and peak co-localization (fraction of BFC peaks falling in channel high regions; Figure 4D). $N_{\text{eq}}$, RMSF are MD-derived; $S_{\phi\psi}$ and the interaction-energy fluctuations $\sigma_E$/CV$_E$ were computed from the raw trajectories; pLDDT is a sequence-derived confidence score included as an orthogonal comparator. All geometric/conformational channels ($N_{\text{eq}}$, $S_{\phi\psi}$, pLDDT, RMSF) are enriched at every threshold, whereas energy fluctuations ($\sigma_E$, CV$_E$) are not. The identical test on shuffled BFC profiles shows no significant enrichment at any threshold for any channel, confirming that the observed co-localization is driven by the positional information of BFC rather than by distributional artifacts. Computed on the full ATLAS library (1,938 proteins).

| | | Region overlap | | Peak co-localization | |
|---|---|---|---|---|---|
| Channel | top-$N\%$ | Enrichment | $p$ | Enrichment | $p$ |
| $N_{\text{eq}}$ | 5% | 1.89× | $< 2 \times 10^{-4}$ | 1.56× | $< 2 \times 10^{-4}$ |
| | 10% | 1.65× | $< 2 \times 10^{-4}$ | 1.55× | $< 2 \times 10^{-4}$ |
| | 15% | 1.47× | $< 2 \times 10^{-4}$ | 1.47× | $< 2 \times 10^{-4}$ |
| | 20% | 1.37× | $< 2 \times 10^{-4}$ | 1.43× | $< 2 \times 10^{-4}$ |
| | 25% | 1.31× | $< 2 \times 10^{-4}$ | 1.39× | $< 2 \times 10^{-4}$ |
| $S_{\phi\psi}$ | 5% | 2.27× | $< 2 \times 10^{-4}$ | 1.73× | $< 2 \times 10^{-4}$ |
| | 10% | 1.83× | $< 2 \times 10^{-4}$ | 1.63× | $< 2 \times 10^{-4}$ |
| | 15% | 1.61× | $< 2 \times 10^{-4}$ | 1.55× | $< 2 \times 10^{-4}$ |
| | 20% | 1.48× | $< 2 \times 10^{-4}$ | 1.49× | $< 2 \times 10^{-4}$ |
| | 25% | 1.39× | $< 2 \times 10^{-4}$ | 1.44× | $< 2 \times 10^{-4}$ |
| pLDDT | 5% | 3.25× | $< 2 \times 10^{-4}$ | 2.40× | $< 2 \times 10^{-4}$ |
| | 10% | 2.49× | $< 2 \times 10^{-4}$ | 2.08× | $< 2 \times 10^{-4}$ |
| | 15% | 2.08× | $< 2 \times 10^{-4}$ | 1.87× | $< 2 \times 10^{-4}$ |
| | 20% | 1.83× | $< 2 \times 10^{-4}$ | 1.73× | $< 2 \times 10^{-4}$ |
| | 25% | 1.66× | $< 2 \times 10^{-4}$ | 1.61× | $< 2 \times 10^{-4}$ |
| RMSF | 5% | 2.40× | $< 2 \times 10^{-4}$ | 1.89× | $< 2 \times 10^{-4}$ |
| | 10% | 2.02× | $< 2 \times 10^{-4}$ | 1.79× | $< 2 \times 10^{-4}$ |
| | 15% | 1.78× | $< 2 \times 10^{-4}$ | 1.64× | $< 2 \times 10^{-4}$ |
| | 20% | 1.62× | $< 2 \times 10^{-4}$ | 1.54× | $< 2 \times 10^{-4}$ |
| | 25% | 1.51× | $< 2 \times 10^{-4}$ | 1.48× | $< 2 \times 10^{-4}$ |
| $\sigma_E$ | 5% | 0.93× | 0.99 | 0.85× | 0.99 |
| | 10% | 0.92× | 1.0 | 0.92× | 0.99 |
| | 15% | 0.94× | 1.0 | 0.93× | 1.0 |
| | 20% | 0.93× | 1.0 | 0.94× | 1.0 |
| | 25% | 0.93× | 1.0 | 0.93× | 1.0 |
| CV$_E$ | 5% | 1.01× | 0.32 | 1.03× | 0.27 |
| | 10% | 0.97× | 0.99 | 0.99× | 0.64 |
| | 15% | 0.97× | 1.0 | 1.00× | 0.52 |
| | 20% | 0.98× | 1.0 | 0.99× | 0.64 |
| | 25% | 0.99× | 0.88 | 1.00× | 0.47 |

Table S4: **Probability that a residue switches backbone conformational state in MD, as a function of BFC score (Figure 5B).** Residues pooled across the 1,930 proteins (termini excluded) are binned into equal-frequency BFC deciles; for each bin we report the mean BFC $z$-score, the fraction of residues that undergo a confirmed backbone-state transition ($\alpha/\beta/\alpha_L$, $\geq$ 5-frame dwell filter, counted within each replica), the binomial 95% confidence interval, and the bin size. The switching fraction rises monotonically from the lowest to the highest decile (bottom 10%: 15.9%; top 10%: 30.4%; enrichment 1.91×). The monotonic trend (per-residue BFC vs. switch/no-switch, pooled within-chain permutation, $\mathrm{NPERM} = 5000$) is significant at $p < 2 \times 10^{-4}$ ($z = 58.9$).

| Decile | Mean BFC $z$ | Switching % | 95% CI | $n$ |
|---|---|---|---|---|
| 1 (lowest) | −0.72 | 15.9 | [15.6, 16.3] | 41,350 |
| 2 | −0.49 | 16.3 | [16.0, 16.7] | 41,349 |
| 3 | −0.40 | 15.6 | [15.3, 16.0] | 41,350 |
| 4 | −0.34 | 16.7 | [16.3, 17.0] | 41,349 |
| 5 | −0.29 | 16.8 | [16.4, 17.2] | 41,350 |
| 6 | −0.24 | 18.1 | [17.7, 18.5] | 41,350 |
| 7 | −0.18 | 18.3 | [17.9, 18.6] | 41,349 |
| 8 | −0.08 | 21.6 | [21.2, 22.0] | 41,349 |
| 9 | +0.31 | 24.1 | [23.6, 24.5] | 41,350 |
| 10 (highest) | +2.42 | 30.4 | [29.9, 30.8] | 41,350 |

Table S5: **BFC peaks co-localize with conformational displacement; both inter- and intra-state components are enriched (Figure 5C).** Total per-residue MD fluctuation was partitioned by the law of total variance into an inter-state (between-basin) and an intra-state (within-basin) component, with global conformational states defined from a C$\alpha$-PCA free-energy landscape. Region-overlap enrichment (symmetric Jaccard) of BFC peaks at inter- vs. intra-state high regions is reported over a pooled within-chain permutation null ($\mathrm{NPERM} = 5000$), in the $N = 592$ proteins with cleanly separated deep basins (inter-basin barrier $\geq$ 2.5 kT). Both components are enriched by $\sim 2\times$ at every threshold; BFC does not distinguish between them, consistent with its training target (ensemble RMSF) being an aggregate of all cross-conformer displacement. Across the full multi-basin set the two components are collinear (per-protein median Spearman 0.75) and cannot be separated.

| | Inter-state displacement | | Intra-state fluctuation | |
|---|---|---|---|---|
| top-$N$% | Enrichment | $p$ | Enrichment | $p$ |
| 5% | 2.14× | $< 2 \times 10^{-4}$ | 2.41× | $< 2 \times 10^{-4}$ |
| 10% | 1.82× | $< 2 \times 10^{-4}$ | 2.01× | $< 2 \times 10^{-4}$ |
| 15% | 1.58× | $< 2 \times 10^{-4}$ | 1.78× | $< 2 \times 10^{-4}$ |
| 20% | 1.44× | $< 2 \times 10^{-4}$ | 1.61× | $< 2 \times 10^{-4}$ |
| 25% | 1.36× | $< 2 \times 10^{-4}$ | 1.51× | $< 2 \times 10^{-4}$ |

Table S6: **BFC peaks co-localize with ligand-induced conformational change across apo–holo protein pairs (Figure 5E).** For 118 apo–holo pairs curated from CryptoSite and PocketMiner, the per-residue conformational change from apo to holo was quantified as $1-$(local CAD-score). BFC profiles were predicted from the apo structure and B-factors alone (the holo state was never seen). Enrichment of BFC high regions/peaks at the high-change residues is reported over a pooled within-chain permutation null ($\mathrm{NPERM} = 5000$), across five top-$N$% thresholds, by symmetric-Jaccard region overlap and peak co-localization.

| | Region overlap | | Peak co-localization | |
|---|---|---|---|---|
| top-$N$% | Enrichment | $p$ | Enrichment | $p$ |
| 5% | 3.29× | $< 2 \times 10^{-4}$ | 2.73× | $< 2 \times 10^{-4}$ |
| 10% | 2.45× | $< 2 \times 10^{-4}$ | 2.29× | $< 2 \times 10^{-4}$ |
| 15% | 1.97× | $< 2 \times 10^{-4}$ | 1.98× | $< 2 \times 10^{-4}$ |
| 20% | 1.69× | $< 2 \times 10^{-4}$ | 1.84× | $< 2 \times 10^{-4}$ |
| 25% | 1.52× | $< 2 \times 10^{-4}$ | 1.64× | $< 2 \times 10^{-4}$ |

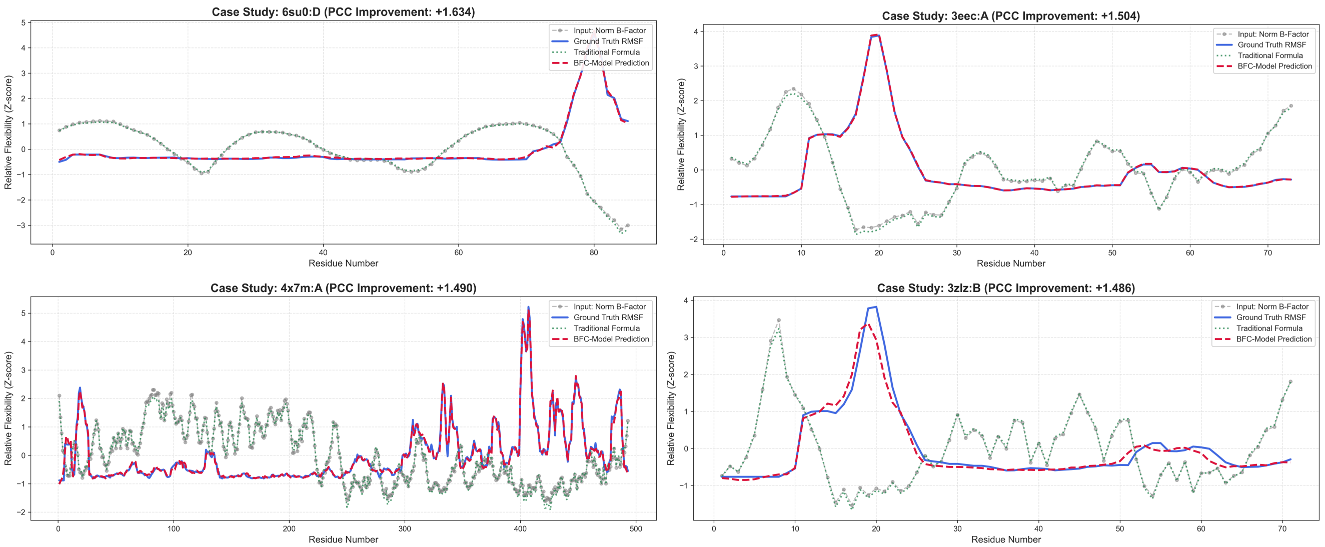


Figure S1: **Extended case studies of dynamic profile reconstruction.** Detailed flexibility predictions for four proteins spanning a range of sequence lengths (6su0, 3eec, 4x7m, and 3zlz). The plots display the BFC prediction (red dashed line) in comparison to the ground-truth RMSF (blue solid line), the input normalized B-factor (gray connected dots), and the traditional formula (green dotted line). The BFC model exhibits consistent alignment with the ground-truth trajectory across all cases, effectively identifying distinct flexibility peaks in both short peptides and larger protein structures such as 4x7m. The PCC improvement scores noted in the panel headers quantify the increase in Pearson correlation coefficient achieved by the model relative to the normalized B-factor input. These metrics validate the capacity of BFC to mitigate experimental artifacts and accurately recover intrinsic biophysical dynamics.

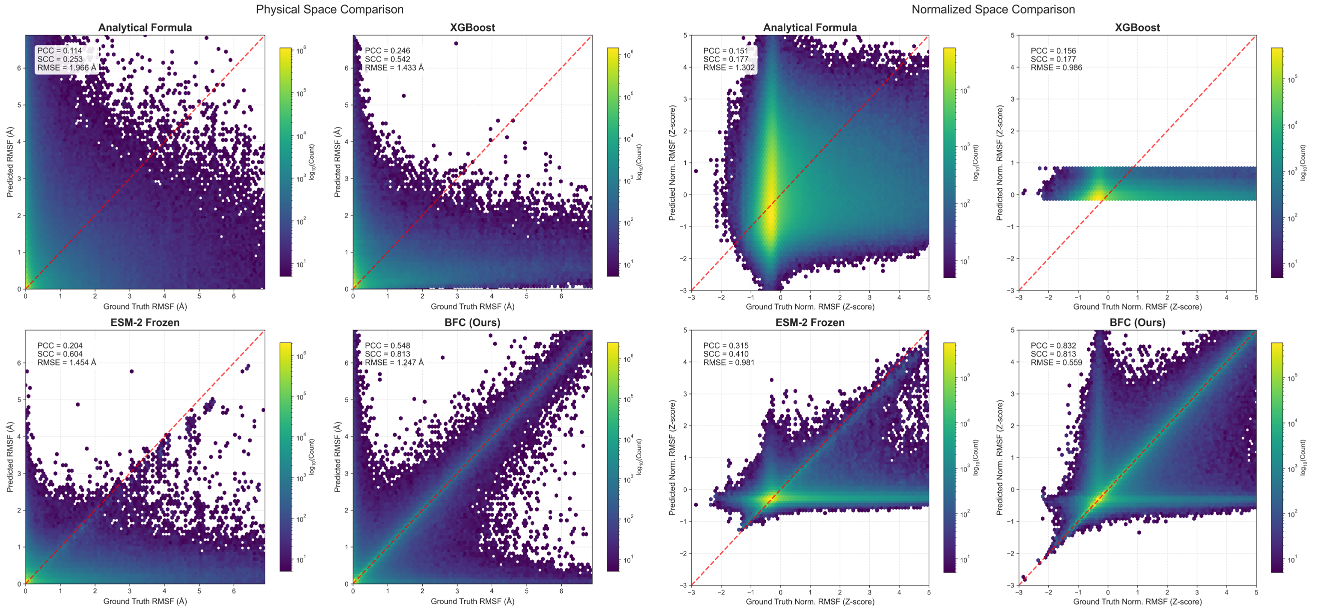


Figure S2: **Comprehensive scatter plot analysis of model performance in physical and normalized spaces.** Density-based comparison of predicted flexibility versus ground-truth RMSF across four methods. The panels are organized into two groups: the Physical Space Comparison on the left measures fluctuations in Angstroms, while the Normalized Space Comparison on the right evaluates performance using Z-scores. The color gradient represents the logarithmic density of data points, where yellow indicates high-concentration regions and purple indicates low density. The red dashed line marks the ideal identity relationship where the prediction equals the ground truth. Baseline methods such as the Analytical Formula and XGBoost exhibit significant dispersion and poor correlation. Notably, in the Physical Space Comparison (left columns), all predictions were subjected to Oracle Rescaling (matched to ground-truth mean/std) to ensure a fair comparison of profile shapes. Despite this, the analytical formula shows broad scatter (SCC=0.25), confirming that B-factors are locally distorted. While the Frozen ESM-2 model shows improved correlation, it still retains considerable variance around the diagonal. In contrast, the BFC model demonstrates superior convergence along the identity line in both evaluation metrics. The statistical indicators provided in each panel confirm that BFC achieves the highest Pearson Correlation Coefficient (PCC) and Spearman Correlation Coefficient (SCC) combined with the lowest Root Mean Square Error (RMSE), validating its robustness across different feature spaces.

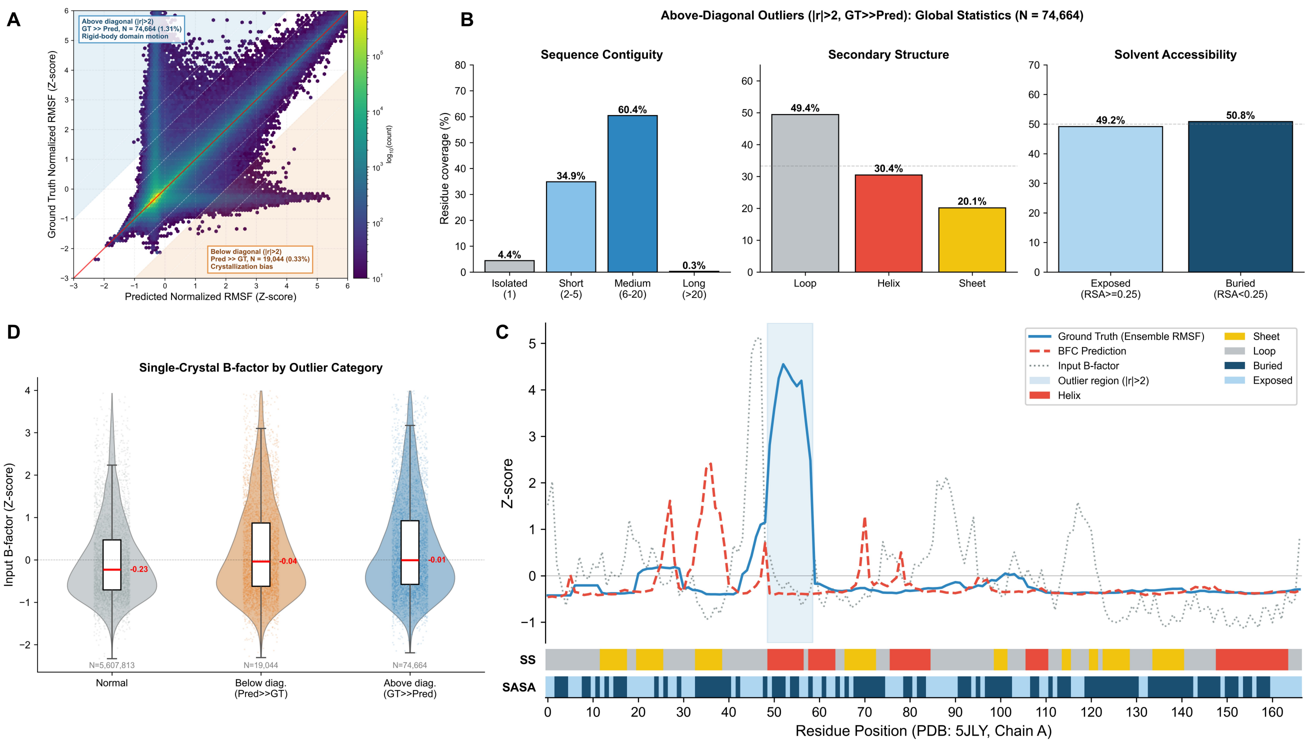


Figure S3: **Diagnostic analysis of prediction outliers resolves the physical origins of discrepancies between model and ground truth.** Outliers are defined as residues exhibiting a prediction residual $|r| = |\mathrm{GT} - \mathrm{Pred}| > 2.0$ in normalized Z-score space. Above-diagonal outliers (GT $\gg$ Pred; N = 74,664, 1.31%) indicate regions where the structural ensemble exhibits high conformational variance that BFC does not attribute to intrinsic local flexibility. Below-diagonal outliers (Pred $\gg$ GT; N = 19,044, 0.33%) indicate latent sequence-encoded flexibility not realized in the experimental ensemble. **A,** Density scatter plot of BFC predictions versus ground-truth ensemble RMSF on the test set (5,701,521 residues). Dashed white lines indicate constant residual magnitudes ($|r| = 1, 2, 3$). The shaded regions highlight the two outlier populations analyzed in subsequent panels. **B,** Global statistical characterization of above-diagonal outliers. Left: sequence contiguity analysis reveals that 60.4% of these outlier residues appear in contiguous blocks of 6–20 residues, indicating coordinated structural displacements rather than isolated prediction errors. Middle and right: secondary structure and solvent accessibility distributions are uniform (Loop 49.4%, Helix 30.4%, Sheet 20.1%; Exposed 49.2%, Buried 50.8%), demonstrating that these outliers are not associated with any specific structural context. **C,** Representative case study (PDB: 5JLY, Chain A, peroxiredoxin-1, 167 residues). The plot overlays three signals: ground-truth ensemble RMSF (blue), BFC prediction (red dashed), and input single-crystal B-factor (gray dotted). In the shaded outlier region (residues 49–58), a buried $\alpha$-helix exhibits extreme ensemble RMSF ($Z > 3.0$) while both the input B-factor and BFC prediction remain near zero, indicating that each individual crystal structure is locally rigid but different crystal forms capture the helix at different spatial positions due to rigid-body domain motion, consistent with the well-documented conformational transitions of peroxiredoxins (Karplus, 2015). **D,** Input B-factor distributions stratified by outlier category. Above-diagonal outliers exhibit near-zero median input B-factors despite their extreme ensemble RMSF, confirming that the high conformational variance originates from inter-crystal positional differences (rigid-body displacement) rather than intra-crystal thermal fluctuation.

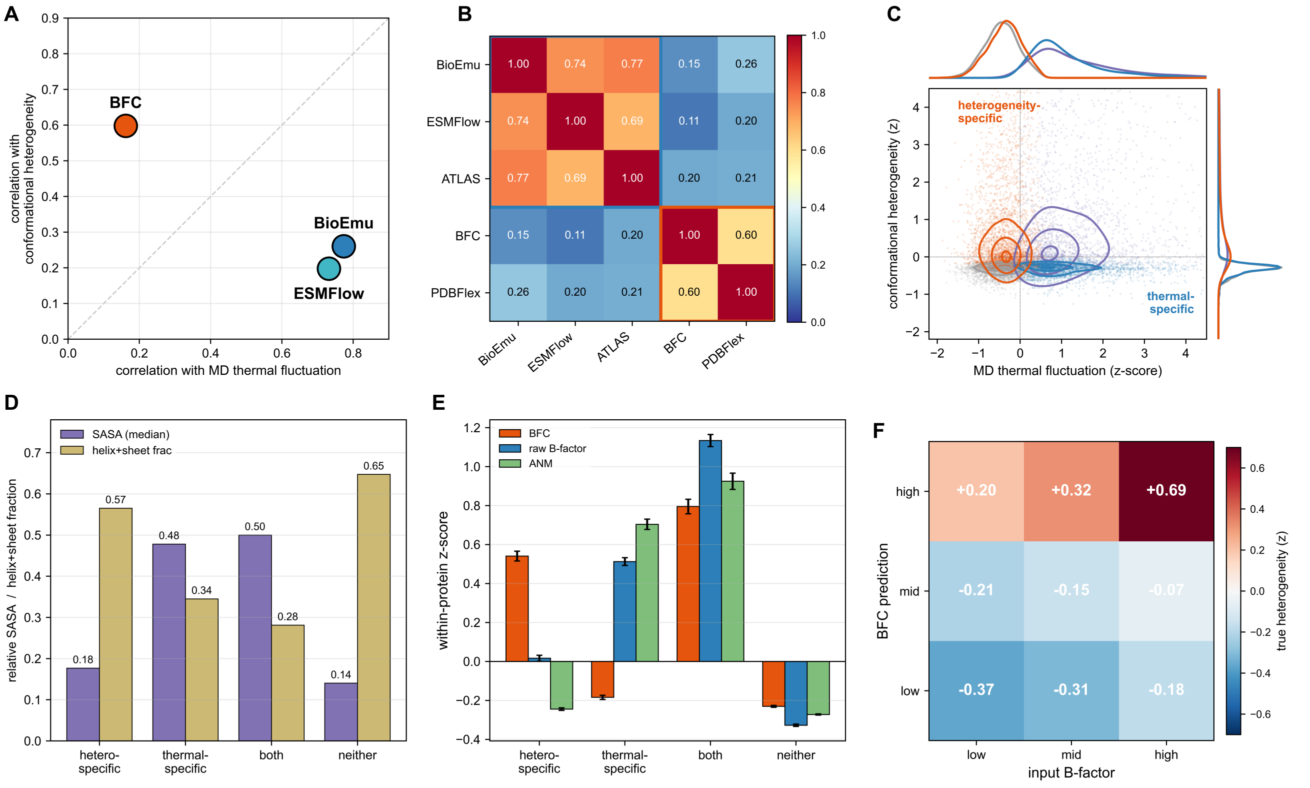


Figure S4: **BFC and ensemble generators capture partially dissociable observables of protein conformational variability.** We generated conformational ensembles for 300 held-out proteins. Of these, 87 lie in the intersection of our test set and ATLAS and so carry both MD-derived RMSF and PDBFlex ensemble-RMSF ground truth, while the remaining 213 were sampled at random from ATLAS. Panels **B–F** require the PDBFlex reference and use only these 87 proteins, whereas panel **A** evaluates the MD-fluctuation axis on all 300 proteins and the heterogeneity axis on the 87. Throughout, ESMFlow denotes the PDB-trained variant (ESMFlow-PDB), fitted to cross-deposition structural heterogeneity rather than to MD trajectories (see Methods). **A,** Each method positioned by its per-protein median Spearman correlation with MD thermal fluctuation (ATLAS-RMSF, $x$, $N = 300$) versus cross-deposition conformational heterogeneity (PDBFlex ensemble RMSF, $y$, $N = 87$). BioEmu (0.77, 0.26) and ESMFlow (0.73, 0.20) align with the MD axis, while BFC (0.16, 0.60) aligns with the heterogeneity axis. **B,** Pairwise per-protein median Spearman matrix ($N = 87$): an MD-fluctuation block (BioEmu/ESMFlow/ATLAS, 0.69–0.77) and a heterogeneity block (BFC/PDBFlex, 0.60) joined by weak cross-terms (0.11–0.26). **C,** Joint density of residues on the two axes (within-chain $z$), with marginal distributions; a subpopulation is heterogeneous across depositions yet low in MD fluctuation (upper-left, *heterogeneity-specific*, 16.2%) alongside a complementary *thermal-specific* set (lower-right, 15.7%). **D,** Structural context (medians): heterogeneity-specific residues are buried (relative SASA 0.18) and secondary-structured (57% helix/sheet); thermal-specific residues are exposed (0.48) loops (66%). **E,** Within-protein $z$-scored scores per residue class: on heterogeneity-specific residues BFC is high (+0.54) whereas experimental B-factor (+0.02) and ANM ($-0.24$) are not. Per-protein paired Wilcoxon (heterogeneity-specific vs. thermal-specific): BFC $p = 4.7 \times 10^{-12}$ (positive), B-factor $p = 5.3 \times 10^{-9}$ and ANM $p = 7.2 \times 10^{-16}$ (both reversed); $N = 87$ proteins. **F,** On strictly held-out test proteins, residues binned into tertiles by input B-factor ($x$) and BFC prediction ($y$), both within-protein $z$; each cell shows the mean ground-truth heterogeneity (PDBFlex $z$; cell counts 1,682–2,990). Ground truth tracks the BFC axis: where BFC's prediction departs from its input B-factor, the held-out PDBFlex reference agrees with BFC (low-B/high-BFC cell +0.20, 95% CI $[+0.06, +0.37]$; high-B/low-BFC cell $-0.18$, $[-0.28, -0.07]$; protein-block bootstrap, $B = 5{,}000$). *Note:* the BFC–PDBFlex value reported here (0.60) is the per-protein median Spearman on this 87-protein subset and differs from the pooled global PCC (0.83, Figure 1, Table S1) because the per-protein metric removes between-chain amplitude variance and the subset spans a restricted dynamic range.

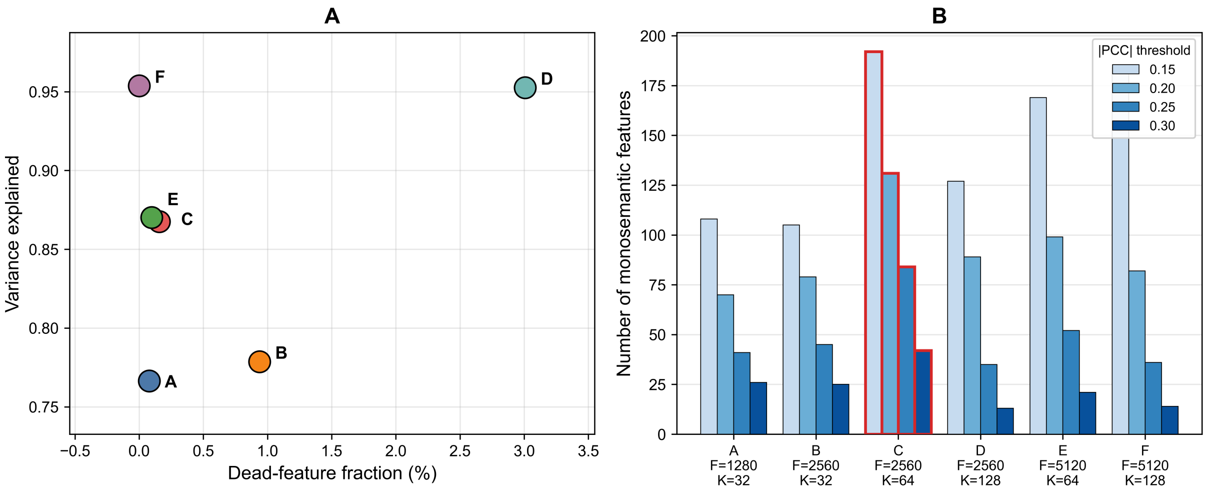


Figure S5: **Selection of the sparse autoencoder configuration.** Six SAE configurations were trained on BFC layer-6 representations and evaluated over the full library ($5.68 \times 10^7$ residues), labelled A–F by their (dictionary size $F$, sparsity $K$) setting: A $(1280, 32)$, B $(2560, 32)$, C $(2560, 64)$, D $(2560, 128)$, E $(5120, 64)$, and F $(5120, 128)$. **A,** Reconstruction–sparsity trade-off: variance explained versus dead-feature fraction. Increasing $F$ and $K$ (configs D, F) pushes variance explained toward $0.95$, but this gain reflects an over-complete dictionary that distributes information across many entangled units rather than improved interpretability. **B,** Number of monosemantic features, features whose activation correlates with ensemble RMSF above a given threshold ($|\mathrm{PCC}|$), counted at four thresholds ($0.15$, $0.20$, $0.25$, $0.30$; red outline marks the selected configuration C). Although the high-capacity configurations achieve marginally higher variance explained, configuration C ($F = 2560$, $K = 64$) consistently yields the largest number of heterogeneity-correlated features at every threshold, indicating that it decomposes the representation into the most interpretable dictionary while keeping reconstruction high (variance explained $0.87$) and dead features negligible ($0.2\%$). C was therefore selected for all subsequent analyses, including the dictionary characterized in Fig. 8.

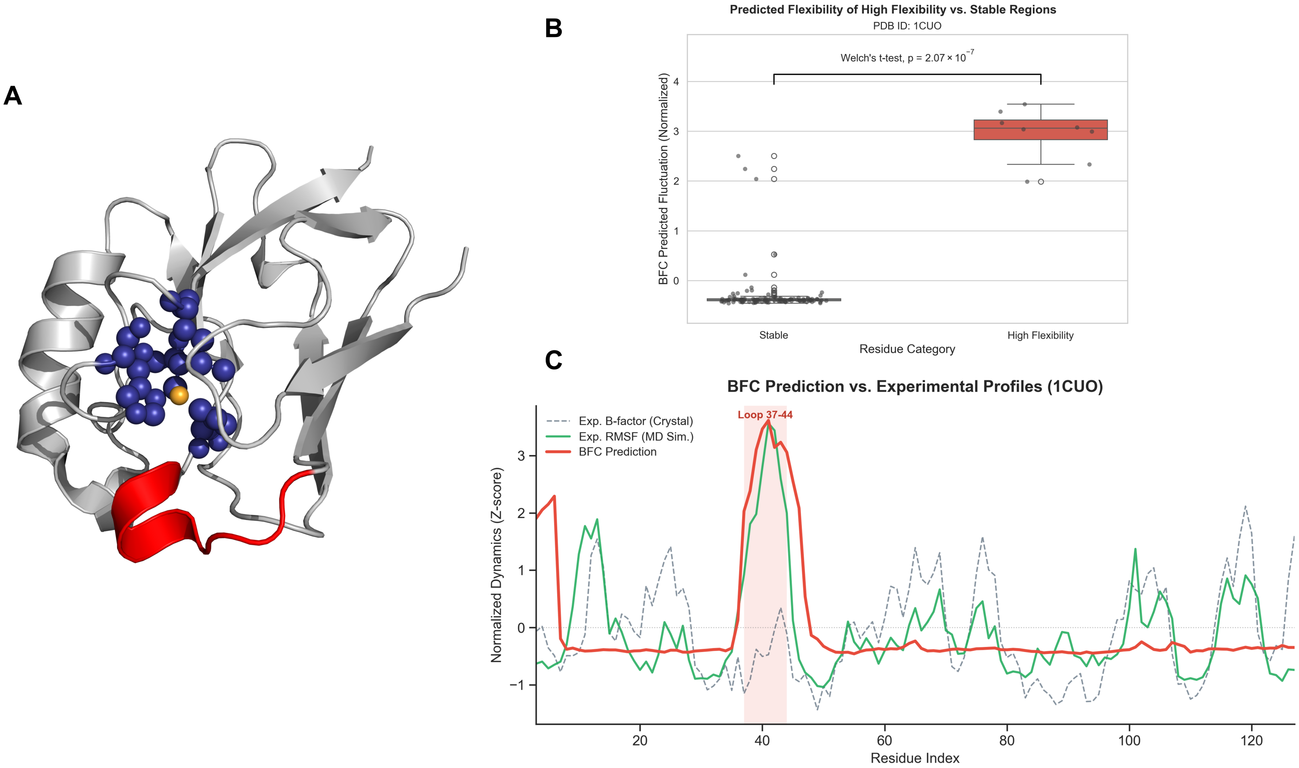


Figure S6: **BFC decodes the functional duality of rigidity and flexibility in Azurin (PDB: 1CUO). A,** Structural visualization of the Azurin active site. The landscape reveals a dynamic duality: BFC correctly assigns structural rigidity to the core residues (blue spheres) that form a precise scaffold to coordinate the copper ion (orange). Simultaneously, the model identifies a distinct, highly flexible loop (residues 37-44, highlighted in red) adjacent to the core whose flexibility has been implicated in forming the intermolecular electron-transfer complex (Inoue et al., 2003). **B,** Statistical validation. A quantitative comparison shows that the BFC-predicted flexibility for the functional loop (red box) is significantly higher than that of the protein's stable regions (Welch's $t$-test, $p = 2.07 \times 10^{-7}$). **C,** Comparative flexibility profiling against external molecular dynamics simulations. The plot overlays the BFC prediction (red solid line) with experimental crystallographic B-factors (gray dashed line) and ground-truth RMSF derived from the ATLAS MD database (green solid line). Notably, while the crystallographic B-factors suggest a dampened motion for the functional loop (residues 37-44) likely due to lattice constraints, BFC accurately recovers the high-flexibility peak consistent with the solution-state MD trajectory. This validates the ability of the model to generalize from structural ensembles to time-resolved simulations.

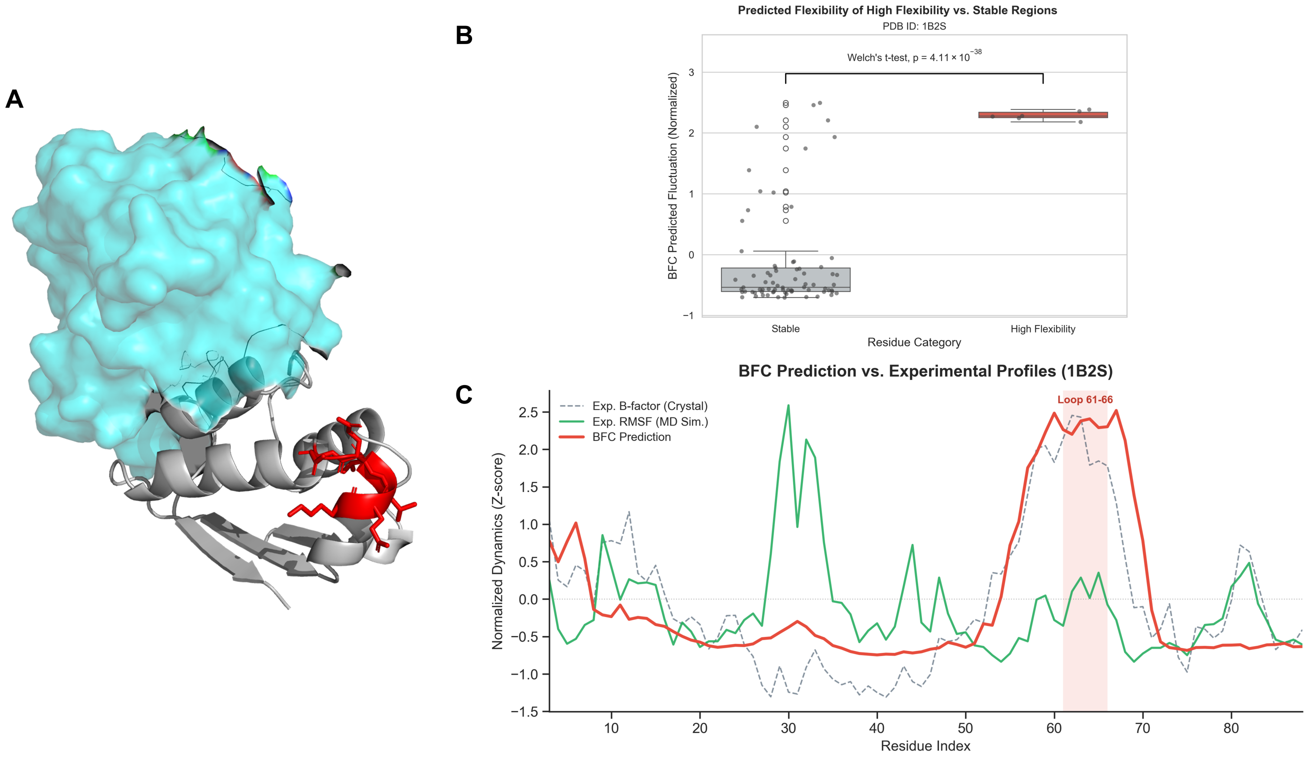


Figure S7: **BFC reveals intrinsic flexibility in Barstar (PDB: 1B2S) consistent with crystallographic evidence. A,** Structural context of the Barstar (Chain E, gray ribbon) and Barnase (cyan surface) complex. The BFC-predicted high-flexibility region (residues 61-66) is highlighted in red, identifying a solvent-exposed loop with high propensity for intrinsic motion. **B,** Statistical validation of the prediction. The BFC-predicted fluctuation scores for the identified loop (red box) are significantly higher than those of the stable scaffold (Welch's $t$-test, $p = 4.11 \times 10^{-38}$). **C,** Comparative flexibility profiling against external molecular dynamics simulations. The plot overlays the BFC prediction (red solid line) with experimental crystallographic B-factors (gray dashed line) and ground-truth RMSF derived from the ATLAS MD database (green solid line). In the loop region (residues 61-66, shaded red), BFC predicts a high-flexibility peak that closely aligns with the experimental B-factors, consistent with the pronounced conformational mobility of this solvent-exposed motif. In contrast, the ATLAS MD simulation exhibits lower fluctuations. We interpret this deviation as reflecting that BFC is trained on structural ensembles that capture the full conformational propensity often reflected in B-factors, whereas individual MD trajectories may sample only a restricted set of metastable states. The result validates that BFC effectively recovers sequence-encoded flexibility consistent with experimental observation.

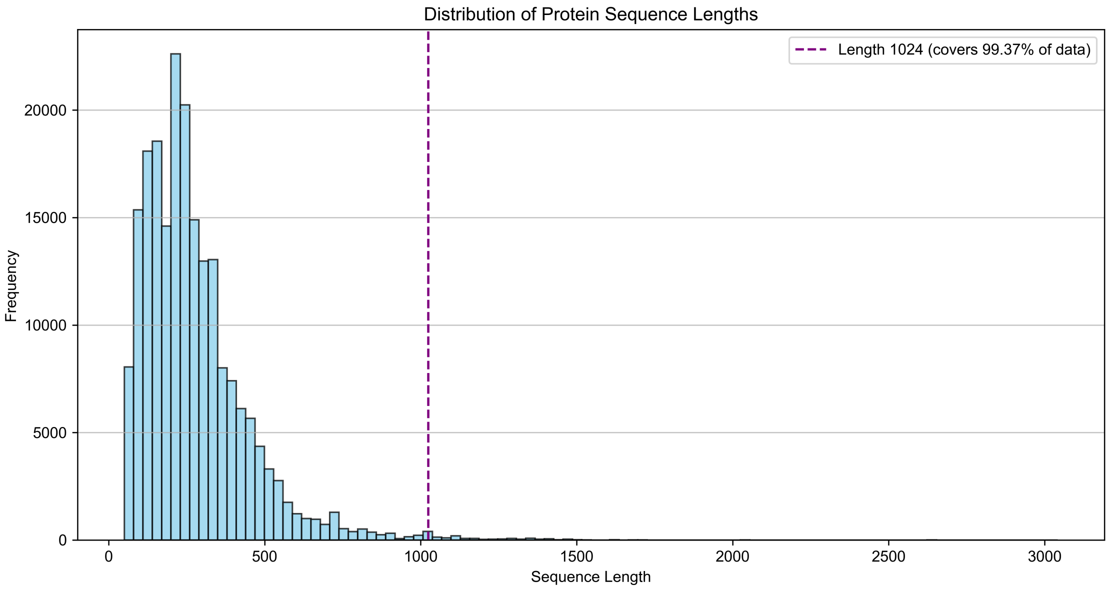


Figure S8: **Distribution of protein sequence lengths.** The vertical dashed line marks the truncation threshold of 1,024 residues set for training and testing. This cutoff covers 99.37% of the training set (206,168 out of 207,465 sequences have a length $\leq$ 1,024), effectively reducing computational overhead while preserving the vast majority of the data.
